# Integrated Cross-Disease Atlas of Human And Mouse Astrocytes Reveals Heterogeneity and Conservation of Astrocyte Subtypes in Neurodegeneration

**DOI:** 10.1101/2025.02.12.637903

**Authors:** Tawaun A. Lucas, Gloriia Novikova, Sadhna Rao, Yuanyuan Wang, Benjamin I. Laufer, Shristi. Pandey, Michelle. G. Webb, Nikolas. Jorstad, Brad A. Friedman, Jesse E. Hanson, Joshua S. Kaminker

**Author notes:** Authors contributed equally to this work (Tawaun A. Lucas, Gloriia Novikova).

## Abstract

Astrocytes play a pivotal role in central nervous system homeostasis and neuroinflammation. Despite advancements in single-cell analyses, the heterogeneity of reactive astrocytes in neurodegenerative diseases, particularly across species, remains understudied. Here, we present an integrated atlas of 187,000 astrocytes from mouse models of Alzheimer’s (AD) and multiple sclerosis (MS) alongside 438,000 astrocytes from AD, MS, and Parkinson’s (PD) patients. Our analysis identified four distinct mouse astrocyte clusters, including two disease-associated astrocyte (DAA) clusters, DAA1 and DAA2. DAA1 displayed reactivity resembling responses to acute stimuli, including endotoxemia, while DAA2 expressed well-known AD risk genes. In an AD model, DAA1 and DAA2 exhibited distinct spatial relationships to amyloid plaques. In humans, we identified eight distinct astrocyte clusters, encompassing homeostatic and disease-associated subtypes. Cross-species analysis linked disease-associated clusters while also highlighting divergent expression in others. Our astrocyte atlas is available through a user-friendly, searchable website: http://research-pub.gene.com/AstroAtlas/.

## Introduction

Astrocytes are one of the most abundant cell types in the brain and play critical roles in maintaining neuronal and synaptic homeostasis. They can become reactive in response to insults or cues they receive from their environment, such as during ischemic stroke^1,2^, after cytokine release from neighboring cell populations^3^, or in response to neuropathologies in neurodegenerative disease^4–6^. Prominent features of reactive astrocytes include cellular hypertrophy and upregulation of proteins, such as GFAP, although these features alone are insufficient to define astrocyte reactivity^7–9^.

Reactive astrocytes can mediate both protective and toxic responses to nervous system injury. For example, reactive astrocytes form glial scars that serve neuroprotective functions and promote recovery from damage^10^. In contrast, reactive astrocytes can also elicit neurotoxic effects by releasing toxic factors such as saturated lipids, causing damage to neurons and oligodendrocytes^3,10,11^. Astrocytes can also lose homeostatic functions, such as when their ability to clear excess glutamate is decreased, resulting in excitotoxicity^12^. The various functional changes observed in reactive astrocytes indicate complex alterations in their transcriptional profiles in response to environmental stimuli that are crucial to understand to decipher the roles of astrocytes in health and disease. Early studies designed to characterize reactive astrocytes using bulk transcriptomic analyses described two states induced by acute traumatic insults in models of ischemic stroke and endotoxemia^1^. However, more recent single-cell RNA sequencing studies in mouse models of chronic neurodegeneration have revealed complex gene expression profiles within the broad population of reactive astrocytes^9,13–18^. Although these recent analyses highlighted transcriptional profiles of reactive astrocytes, a more detailed description of the heterogeneity of reactive astrocytes across chronic neurodegenerative diseases, including Alzheimer’s Disease (AD) and Multiple Sclerosis (MS), is lacking.

Additionally, while integrative analyses of microglia and oligodendrocytes have successfully identified distinct subpopulations of disease-associated cells^19–25^, a similar comprehensive analysis has not been conducted for astrocytes, limiting our understanding of astrocyte populations that may be relevant to disease.

The recent explosive increase of single-cell and single-nuclei RNA sequencing datasets from human patient brains and rodent models provides an opportunity to leverage these data to perform a broad and detailed characterization of astrocytes, providing a better understanding of the diversity and complexity of their transcriptional programs. Therefore, we conducted an integrative meta-analysis using mouse models of AD and MS and human patient astrocytes from AD, MS, and Parkinson’s Disease (PD) samples. Our analysis included 187,000 mouse astrocytes from 181 samples derived from 6 different AD and 3 MS models. Additionally, we analyzed 438,000 human astrocytes from 6 AD, 4 MS, and 3 PD studies. Importantly, we computationally removed ambient RNA from all data sets before integration to more accurately characterize astrocytes and to help identify underrepresented cell states^26–28^. Our integrated mouse analysis revealed two distinct disease-associated populations of astrocytes found within both AD and MS models. Comparison to previously generated data sets and additional data generated for this study suggests that these distinct populations correspond to 1) a reactive state that is also found in acute disease models (e.g. endotoxemia) and 2) a reactive state that is more unique to neurodegeneration models. Importantly, we show that these two populations have distinct spatial distribution patterns relative to disease pathology in an AD model. Our integrated human analysis identified previously uncharacterized and distinct disease-relevant clusters in MS and AD patients. We performed a cross-species analysis and identified those mouse disease-associated clusters that best correspond to human disease-relevant clusters, which can provide insight into the relevance of different models for modeling disease biology and developing therapeutics. Together, our data provide a comprehensive description of astrocytes across both mouse models and human neurodegenerative disease tissue samples, providing unprecedented resolution of the transcriptional landscape of astrocytes in neurodegenerative disease.

## Results

### Development of an integrated single-cell atlas of astrocytes in neurodegenerative disease

To develop an integrated single-cell atlas, we collected a broad set of publicly available MS and AD mouse models and human AD, MS, and PD patient astrocyte data **(Figure 1A and Table 1 & <u>2</u>)**. Prior to integration, we applied rigorous pre-processing steps to each data set **(Figure 1A and Methods)**. In particular, our workflow aligns all reads to the same genome, calls cells using *CellRanger*, removes ambient RNA contamination using *CellBender*^28^, filters doublets using *scDblFinder*^29^, and annotates coarse cell types using label transfer from the same reference dataset^30^. Likely non-astrocytic cells were identified by scoring cells in the initial integrated space using previously established brain cell type-specific markers (see Methods), and high-scoring clusters were removed. Our approach generated a comprehensive compendium of high-quality astrocytes from disease and normal tissue from both mouse and human samples.

**Figure 1:**
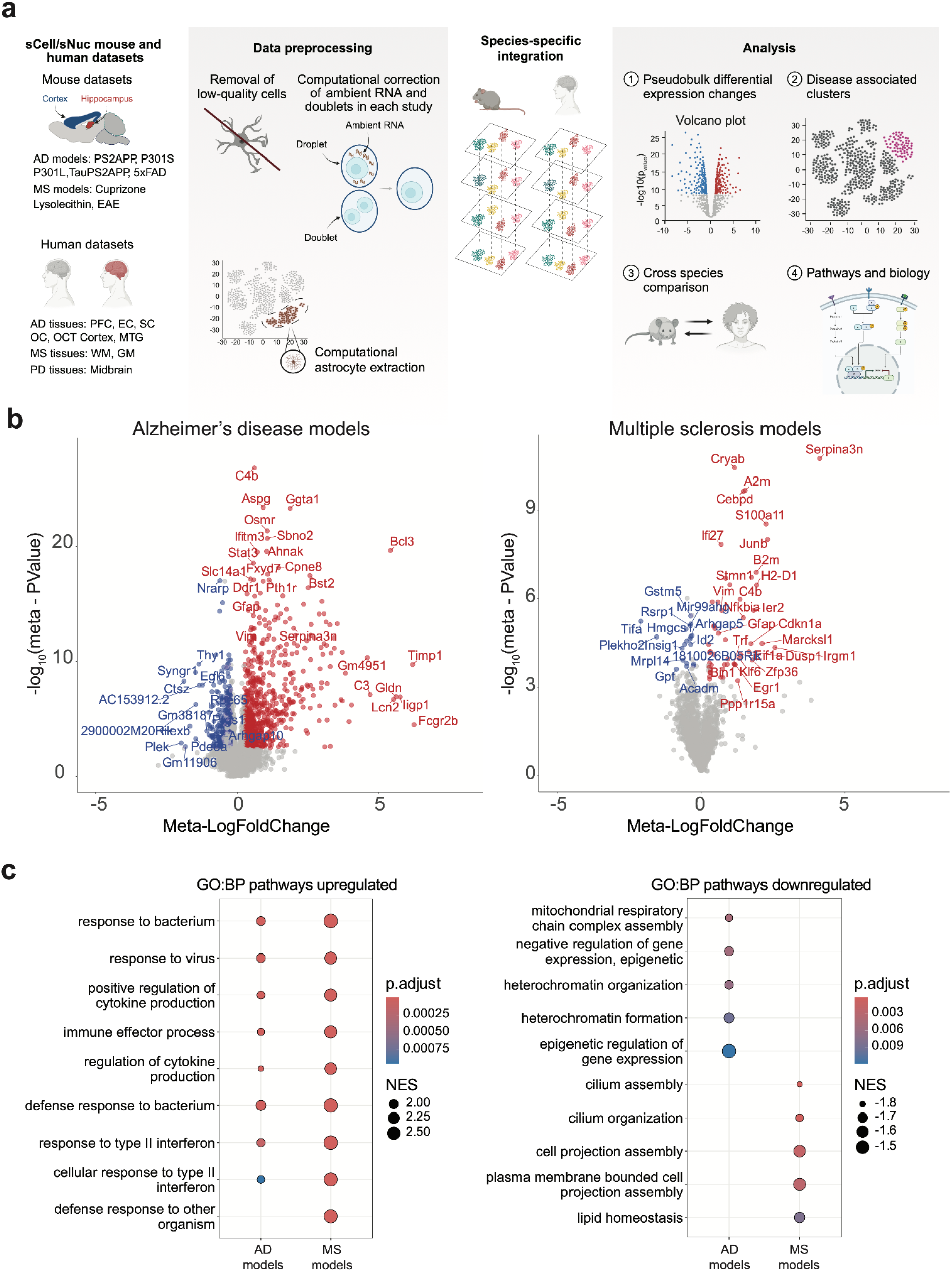
Integration of Mouse Neurodegenerative Datasets reveals disease-specific changes in astrocytes. (A) Schematic depicting computational analysis pipeline highlighting pre-processing steps. (B) Volcano Plot indicates differential expression between disease and normal astrocytes, Alzheimer’s disease models (left), and Multiple Sclerosis models (right). Red and blue colors indicate significantly differentially expressed genes (|meta-logfoldChange| > 0.5, metaFDR < 0.05), and grey indicates no significant change. (C) Gene Set Enrichment Analysis of Gene Ontology Biological Pathways upregulated in AD models or MS models (left) or downregulated in AD models or MS models (right). The color of the circles indicates the adjusted p-value, and the size indicates the Normalized Enrichment score (NES) for the specific pathway. Row names include the top 5 scoring gene ontology categories for each disease.

**Table 1.** Table of datasets included in the mouse astrocyte atlas.

| GEO Accession | First Author | Last Author | Brain Region | Disease Model | Model | Sample Size | # astrocytes | Data type | Library Prep |
| --- | --- | --- | --- | --- | --- | --- | --- | --- | --- |
| GSE140511 | Zhou, S. | Colonna, M. | Left Hemisphere | AD | 5xFAD | 6 (6M,0F) | 9,075 | Nuclei | 10x |
| GSE143758 | Habib, N. | Schwartz, M. | Hippocampus | AD | 5xFAD | 30 | 24,689 | Nuclei | 10x |
| GSE166261 | Shi, Y. | Holtzman, DM. | Hippocampus | AD | P301S | 6 (pooled animals) | 735 | Nuclei | 10x |
| GSE161227 | Zhao, N. | Ren, Y. | Cortex | AD | 5xFAD | 24 (12M,12F) | 24,958 | Whole-cell | 10x V3 |
| GSE161224 | Zhao, N. | Ren, Y. | Cortex | AD | 5xFAD | 24 (12M,12F) | 46,674 | Whole-cell | 10x V3 |
| GSE150934 | Choi, H. | Lee, DS. | Hippocampus | AD | 5xFAD | 4 | 449 | Whole-cell | 10x |
| GSE153895 | Lee, SY | Hansen, D. | Hippocampus | AD | PS2APP, P301L | 12(M) | 2,870 | Whole-cell | 10x |
| GSE181786 | Lee, SY | Bohlen CJ | Hippocampus | AD | PS2APP, TauPS2APP | 11(M) | 6,728 | Whole-cell | 10x |
| GSE160512 | Lee, SY | Bohlen, CJ | Hippocampus | AD | PSAPP | 12(F) | 3,093 | Whole-cell | 10x |
| GSE180041 | Wang Y. | Hanson,J | Hippocampus | AD | TauP301s | 9 | 13,773 | Whole-cell | 10x V2 |
| GSE148676 | Shen, K. | Friedman, B. | Corpus Callosum | MS | Cuprizone | 20 | 2,647 | Whole-cell | 10x V2 |
| GSE129763 | Wheeler, MA | Li, Z. | Whole brain (sorted) | MS | EAE | 12 | 12,186 | Whole-cell | drop-seq |
| GSE129609 | Wheeler, MA | Li, Z. | Whole brain | MS | EAE | 27 | 6,891 | Whole-cell | drop-seq |
| GSE182846 | Shen Y. | Freidman, B. | Corpus Callosum | MS | Lysolecithin | 12 | 683 | Whole-cell | 10x V2 |

Our initial analysis focused on mouse astrocytes. We integrated data from astrocytes isolated from different AD and MS mouse models and their respective controls, including data from 12 publicly available datasets comprising 181 samples and 205,000 astrocytes. After rigorous QC, we included 187,000 astrocytes in our integrative analysis **(Table 1)**. Among the AD models, our integrated data set contains amyloidosis-only models (PS2APP mice^31,32^, 5xFAD mice^33,34^), tauopathy models (TauP301S^35^ and TauP301L^36,37^), and TauPS2APP animals with both amyloid and tau pathology^30,37,38^. Among the MS models, we included astrocytes from mice that underwent cuprizone- or lysolecithin-induced demyelination as well as astrocytes from mice with experimental autoimmune encephalitis (EAE) induced by injecting MOG_35–55_^39,40^. Datasets from AD models were generated from either the hippocampus or cortex, datasets from demyelination models were generated from the corpus callosum, and datasets from the EAE model were generated from the whole brain. The number of astrocytes isolated per sample varied by study **(Table 1)**.

### Differential expression across disease models highlights common and distinct features of astrocytes in AD and MS models

As our integrated atlas of mouse astrocytes reflects responses to a wide range of pathologies from distinct pre-clinical models, we first asked if astrocytes from AD or MS models shared common and/or distinct gene expression changes. For this, we performed a pseudobulk analysis comparing astrocytes from disease models with astrocytes from their respective matched controls. To identify robust gene expression changes consistent across multiple datasets, we employed a meta-analysis approach^41^. This analysis highlighted significant and robust alterations in gene expression related to astrocyte reactivity in both the AD and MS models. For example, marked increases in the expression of *C4b*, *Serpina3n*, Gfap, and *Vim* **(Figure 1B; Supplemental Table 1)** were observed across both AD and MS models. We also used each model’s ranked differentially expressed genes (DEGs) to conduct gene set enrichment analysis (GSEA; see Methods). GSEA of data from both the MS and AD models indicated a pronounced enrichment of immune regulatory responses, including responses to interferon/bacterium and regulation of cytokine production **(Figure 1C; Supplemental Table 1)**. These results suggest that astrocytes from the AD and MS models potentially share a core transcriptional response between the distinct disease states involved in immune regulatory functions. Astrocytes are crucial for responding to neuroinflammatory signals and maintaining neuronal health, and this may necessitate a conserved response mechanism across different neurodegenerative conditions. In contrast to the analysis of the upregulated genes, the pathways revealed by the downregulated genes from AD or MS models were quite distinct, which may reflect different aspects of astrocyte function between disease models. The top pathways downregulated in AD models were mitochondrial respiratory chain complex, heterochromatin organization, and negative regulation of gene expression. The downregulation of these pathways suggests impaired energy production, which could compromise the astrocytic support of neurons and exacerbate reactive and neuroinflammatory responses. The top pathways downregulated in MS models were related to cilium assembly and organization.

Astrocyte cilia are involved in regulating astrocyte morphology and neurodevelopment^42^ and inflammatory responses, and disruption in their assembly could exacerbate the inflammatory environment or hinder the repair and maintenance of tissue, contributing to the inflammatory-induced neurodegeneration seen in MS^43^.

### Integration of astrocytes from mouse models identifies 4 distinct astrocyte gene programs, including two disease-associated clusters

To characterize astrocytes at the level of single-cell gene expression, we used *Canonical Correlation Analysis*^44^ to integrate all of the mouse datasets and identify shared transcriptional profiles of astrocytes. Astrocytes were clustered using an iterative approach, ensuring that each cluster had at least 5 unique marker genes compared to other astrocyte clusters (see Methods).

This approach yielded an integrated space containing four high-quality subpopulations of astrocytes with an even distribution of cells from each study **(Figure 2 A-C),** providing confidence in our integration approach.

**Figure 2:**
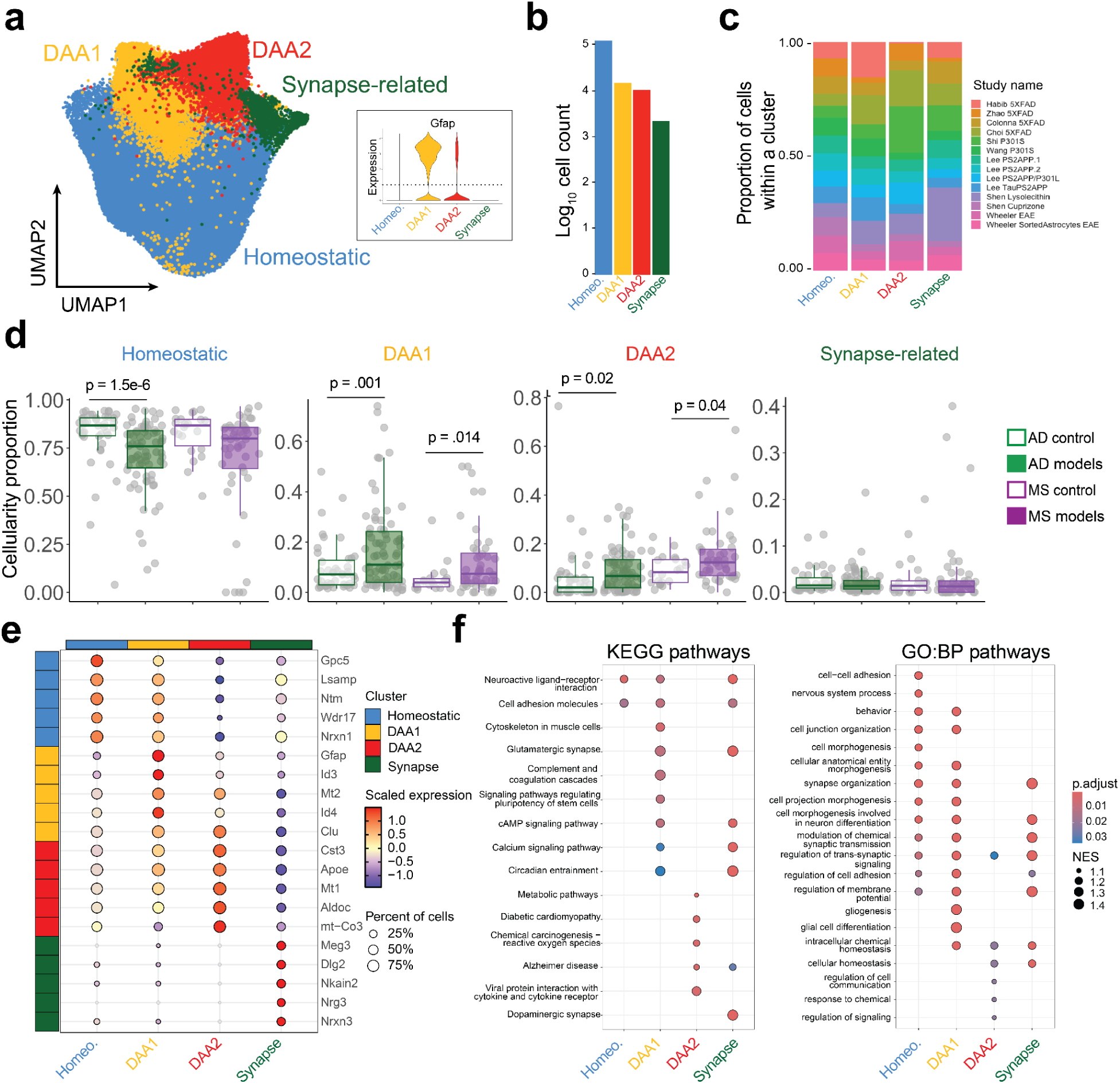
**Cell-level clustering of mouse astrocytes reveals multiple disease-associated astrocyte clusters.** (A) Uniform Manifold Approximation and Projection (UMAP) of astrocytes from mouse models of neurodegeneration and healthy littermates. Iterative clustering (see Methods) resulted in four distinct clusters, and the inset violin plot shows GFAP expression within each cluster. (B) Counts of cells by cluster. (C) Proportion of cells per cluster, colored by study. (D) Barplots indicating cellularity percent by cluster and disease indication. Each dot represents a sample from an individual study. The y-axis represents the sample’s percentage of cells contributing to a specific cluster. P-values are indicated over groups with significant differences in cellularity and were generated from the Kruskal-Wallis test and post-hoc Wilcox test on center-log transformed proportions (see Methods). (E) Dotplot of the top 5 cluster markers of each cluster. Color represents the scaled expression of each cell in that cluster. Dot size represents the percentage of cells that express that specific marker. (F) Dotplot of top differential pathways KEGG (left) and Gene Ontology Biological Process pathways (right). The color of the circles indicates the adjusted p-value, and the size indicates the Normalized Enrichment Score (NES) for the specific pathway. Row names include the gene ontology categories. Pathways were identified using Gene Set Enrichment analysis on Gene Ontology biological pathways.

Of the 4 clusters identified, one cluster contained the largest proportion of cells. We termed this cluster “Homeostatic” as it included a higher proportion of cells from samples of non-transgenic and control conditions **(Figure 2D)**. We termed the smallest cluster “Synapse*-*Related” as astrocytes in this cluster had marker genes related to synapse function, such as *Dlg2* and *Nrxn3* **(Figure 2E)**. Astrocytes in this synapse-related cluster were not differentially abundant between disease and matched control samples **(Figure 2D),** and recent literature also identified a similar cluster of astrocytes expressing synaptic genes ^45,46^. Interestingly, both of the remaining two clusters were differentially abundant between disease samples and matched controls, and as such, were termed “Disease-Associated Astrocyte Type 1” (DAA1) and “Disease-Associated Astrocyte Type 2” (DAA2) **(Figure 2D).** Marker genes for the DAA1 cluster included canonical reactive astrocyte markers such as *Gfap* and *Id3* (**Figure 2E; Supplemental Table 2**). Of note, marker genes for the DAA2 cluster included the AD risk-associated genes *Apoe* and *Clu* **(Figure 2E),** which may suggest that this subpopulation of astrocytes may be critical for understanding the effects of these genes during neurodegeneration. Using ranked DEGs comparing each cluster to the other astrocytes, we performed GSEA analysis on KEGG and GO Biological Process pathways (**Figure 2F**). The Homeostatic and Synapse-related clusters had increased expression of genes associated with pathways, including cell adhesion and synaptic transmission. The enrichment of these pathways was primarily driven by increased expression of neurotrophic adhesion molecule genes such as *Mdga2*, *Grid2*, and *Nrxn1* in the Homeostatic cluster and *Nrg3*, *Dlg2*, and *Ptprd* expression in the Synapse-related cluster. The pathways enriched in the DAA1 cluster included gliogenesis and the complement cascade, influenced by *C4b*, *F3,* and *Serpine2.* In contrast, pathways enriched in the DAA2 cluster were associated with cell communication, driven by *Apoe*, *Dbi*, and *Clu*.

Next, we performed differential expression analysis between DAA1 and DAA2 to understand their transcriptional programs. Our analysis highlighted significant differences in gene expression **(Figure 3A)**, suggesting distinct reactivity states (398 genes up-regulated in DAA1; 280 genes up-regulated in DAA2). GO enrichment analysis between DAA1 and DAA2 revealed that DAA1 is enriched in annotations for *cell junction a*nd *adhesion*, while DAA2 shows enrichment in translation related pathways and aerobic respiration **(Figure 3D)**.

**Figure 3:**
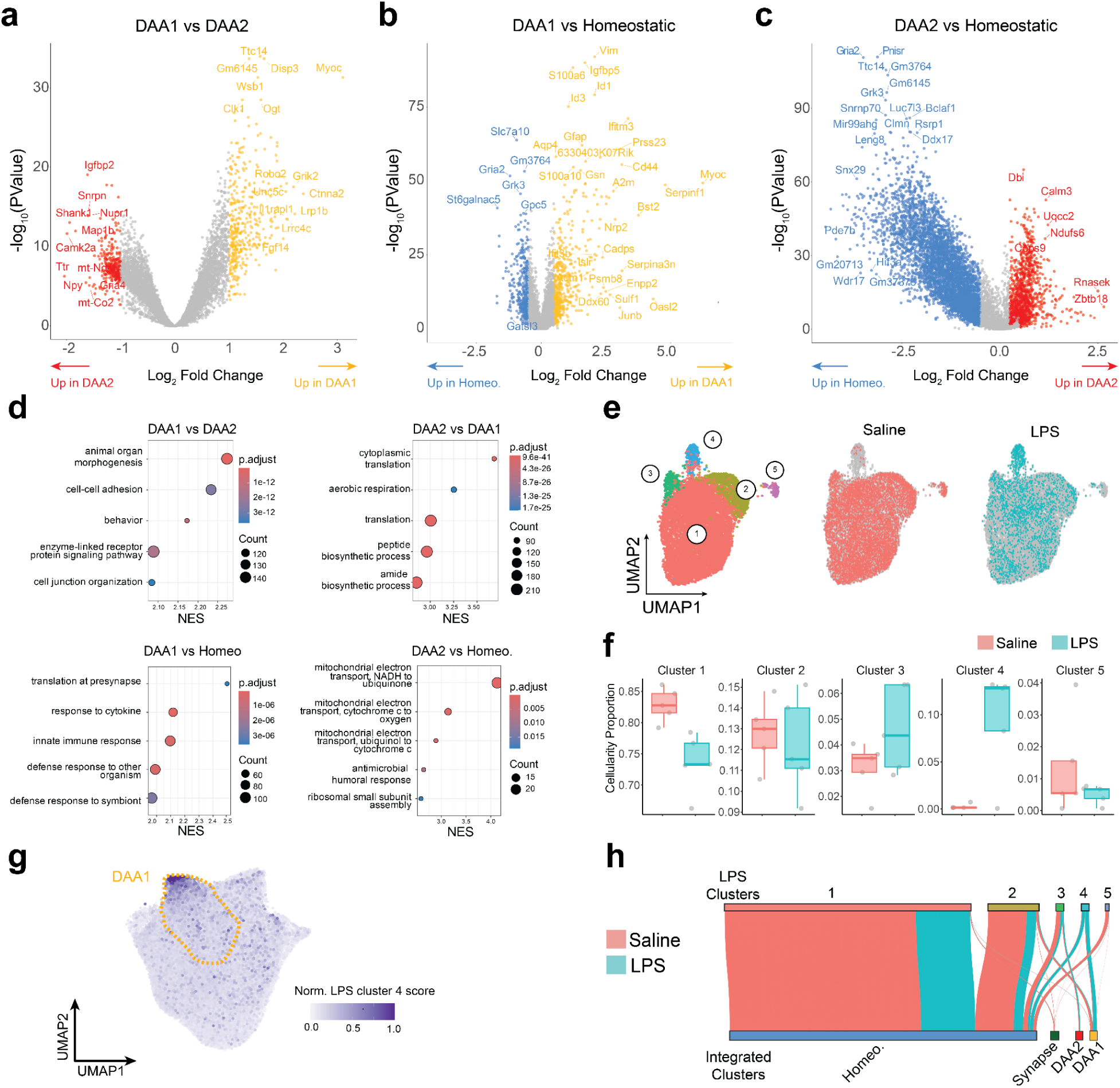
**Differences between DAA1 and DAA2 astrocytes and comparison to reactive astrocytes from acute insult models.** (A) Volcano Plot of PseudoBulk differential expression analysis of DAA1 vs DAA2 astrocytes. Gold indicates significance in DAA1, red indicates significance in DAA2 (|logfoldChange| > 1, p < 0.05), and grey indicates no significant change. (B) Volcano Plot of PseusdoBulk differential expression analysis of DAA1 vs. Homeostatic astrocytes. Gold indicates significance in DAA1, blue indicates significance in Homeostatic ( |logfoldChange| > 1, p < 0.05), and grey indicates no significant change. Volcano Plot of PseudoBulk differential expression analysis of DAA2 vs. Homeostatic. Red indicates significance in DAA2, blue indicates significance in homeostatic ( |logfoldChange| > 1, p < 0.05), and grey indicates no significant change. *(C)* Dotplot of GSEA of Gene Ontology Biological Pathways on differentially expressed genes between DAA1 and DAA2 (top). Top 5 categories are shown for each subtype, as indicated. The color of the circles indicates the adjusted p-value, and the size indicates the ratio of genes in that pathway. GSEA of Gene Ontology Biological Pathways for DAA1 vs Homeostatic, or DAA2 vs Homeostatic(bottom). Top 5 categories for each comparison are shown in dotplot. The color of the circles indicates the adjusted p-value, and the size indicates the Normalized Enrichment Score for the specific pathway. (E) Uniform Manifold Approximation and Projection (UMAP) of mouse LPS stimulated and control astrocytes. Circled numbers indicate cluster numbers (F) Barplots indicating cellularity percent by cluster and treatment. Each dot represents an individual mouse. The y-axis represents the sample’s percentage of cells contributing to a specific cluster. Cluster numbers are indicated at the top of each plot.(G) Scoring integrated neurodegeneration astrocytes shown in Figure 2A with differentially expressed genes from LPS-stimulated astrocytes. (H) Sankey plot of the predicted clusters of cells following cell-level PCA projection of LPS-stimulated astrocytes into our integrated dataset. The sankey plot shows that DAA1 in integrated space, primarily maps onto the LPS-induced cluster 4.

To further explore these differences, we compared each cluster to the Homeostatic cluster **(Figure 3B & C)**. Compared to the Homeostatic population, pathways upregulated in DAA1 were related to cytokine and immune responses and translation at the synapse **(Figure 3D)**. Similarly, comparing DAA2 to homeostatic astrocytes highlighted pathways related to immune response. However, the DAA2 versus Homeostatic comparison also highlighted the upregulation of pathways associated with the mitochondrial electron transport chain, which could reflect cellular respiration alterations seen in AD^47^ and MS^48^.

### Mouse DAA1 signatures resemble signatures derived from astrocytes after acute LPS stimulation

Next, we compared DAA1 and DAA2 populations to reactive astrocytes from acute LPS treatment. For this purpose, we injected mice intraperitoneally with LPS or saline and collected brains 48 hours later for single-cell RNA sequencing analysis of the hippocampus. We processed and subsetted astrocytes using the same protocol as our integrated map, yielding five distinct clusters, including Cluster 4, which increased after LPS treatment **(Figure 3E & F)**.

We performed two analyses comparing astrocytes from LPS-treated mice to our atlas from disease models. The first involved differential expression analysis between LPS-treated and PBS-treated astrocytes, identifying the set of upregulated genes in response to LPS treatment **(Supplemental Figure 3A)**. We then calculated the average expression of this set of 75 LPS-induced genes for each astrocyte in our neurodegeneration atlas, and strikingly, the astrocytes in our atlas that scored highest were those within the DAA1 cluster **(Figure 3G)**. Similarly, the set of upregulated DEGs from a previous bulk LPS study or another acute model of inflammation, the middle cerebral artery occlusion (MCAO) model of acute ischemic stroke^1,49^, also predominantly mapped onto cells within the DAA1 cluster **(Supplemental Figure 3B)**. For the second analysis, we compared cluster identities from the two datasets by projecting cells from our single-cell LPS dataset into the PCA space of our integrated atlas (see Methods). We scored similarity between the nearest neighbors and visualized prediction scores using a Sankey plot **(Figure 3H)**. This analysis highlights that cells from the LPS Cluster 1 predominantly map to the Homeostatic cluster, while cells from the LPS-enriched cluster 4 predominantly map to the DAA1 cluster.

Together, these data indicate that DAA1 astrocytes most closely resemble acutely stimulated astrocytes. While there are many differences between responses to LPS, amyloid, tau, demyelination, and EAE, one interpretation of our analysis is that some reactive astrocytes in chronic disease models may exhibit transcriptional signatures similar to acute inflammatory responses like those to LPS or MCAO (DAA1), while other reactive astrocytes (DAA2) may arise from a distinct, more chronic inflammatory brain environment seen in models of AD and MS.

### Mouse DAA1 and DAA2 astrocytes exhibit distinct spatial distribution relative to amyloid plaques

Next, we wanted to characterize the spatial distribution of DAA1 and DAA2 populations in the context of disease pathology. We used TauPS2APP mice for this analysis and performed *in-situ* hybridization using multiplexed RNAscope and antibody staining to identify astrocyte subtypes and amyloid pathology^50^. The markers we used included DAPI to identify nuclei, *6E10* antibody staining to identify amyloid plaques, *Slc1a3* RNA as a pan-astrocyte marker^51^, and the combination of *Apoe* and *Igfbp5* RNA to discriminate between DAA1, DAA2, or other astrocytes **(Figure 4A, B)**. In particular, among *Slc1a3* expressing cells, DAA1 cells were identified by high *Igfbp5* and moderate *Apoe* expression, and DAA2 cells were identified by low *Igfbp5* and higher *Apoe* expression **(see Methods; Figure 4A, B and Supplemental Figure 3C).** Using this approach, we observed a significant enrichment in the expression of DAA marker genes, and the proportion of DAA populations counted in TauPS2APP animals compared to non-transgenic controls **(Supplemental Figure 3C)**.

**Figure 4.**
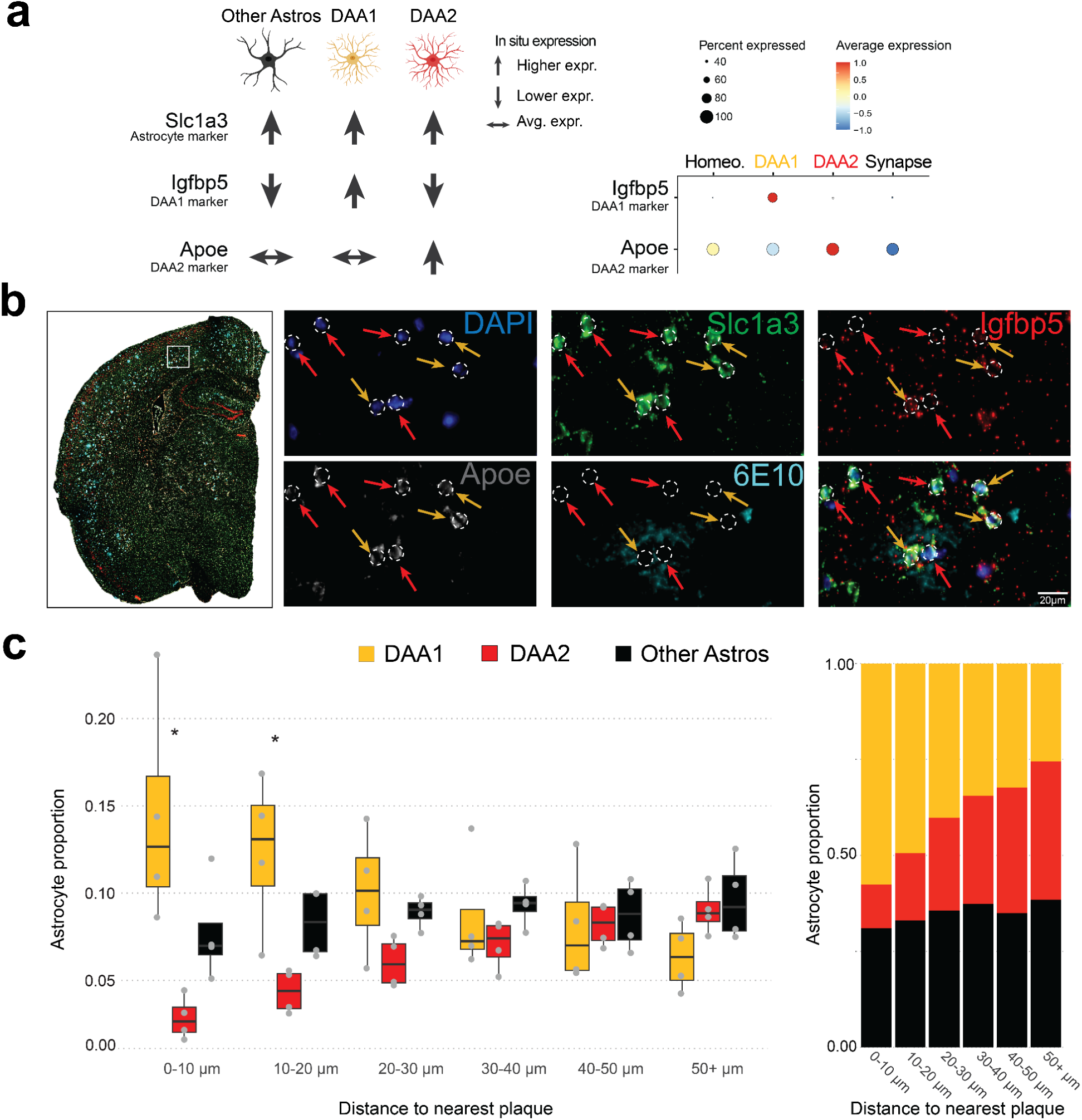
**DAA1 And DAA2 have distinct spatial relationships to amyloid plaques.** (A) Graphical representation of the combination of marker gene expression profiles used to identify astrocyte subtypes with in-situ RNA analysis (left), and dotplot showing expression of those marker genes from the integrative atlas (right). Dot color represents the scaled expression of each cell in that cluster and dot size represents the percentage of cells that express that specific marker. See Supplemental Figure 3C-E for details of image analysis (B) Low-resolution representative photomicrograph of TauPS2APP mouse brain section (left). Boxed region reflects the region shown in high resolution images depicting the relationship of astrocyte subtype markers to amyloid plaques (DAPI in blue; Slc1a3 in green; Igfbpf in red; Apoe in gray; 6e10 in turquoise; scalebar = 20μm). (C) Quantification of cortex and hippocampus astrocyte subtypes by distance to amyloid plaques (n = 4 TauPS2APP mice, average value for all astrocytes across two sections is shown for each mouse). (left) Proportion of astrocyte subtype by distance to nearest plaque depicted as boxplot. Proportion was calculated as the total number of an astrocyte subtype / total astrocytes at a given distance bin from amyloid plaques. Stars represent significant comparisons (Anova with Bonferroni correction;n = 4 mice p <0.001). (right) Ratio of astrocyte subtype by distance to nearest plaque depicted as stacked bar plot.

Next, using this approach, we examined the spatial distribution of DAA1, DAA2, or non-DAA astrocytes by quantifying the proportion of each astrocyte subtype relative to the distance to amyloid plaques. Strikingly, our analysis revealed distinct spatial distributions of DAA1 and DAA2 astrocytes. DAA1 astrocytes were significantly higher in abundance compared to DAA2 astrocytes in the regions closest to amyloid plaques, and the proportion of DAA1 astrocytes decreased progressively with increasing distance from the plaques **(Figure 4C)**. In contrast, DAA2 astrocytes were relatively lower in abundance near amyloid plaques, and their proportion progressively increased as the distance from the plaques increased **(Figure 4C).** This spatial distribution of DAA1 astrocytes, which we found are more transcriptionally similar to acutely reactive astrocyte populations, suggests that DAA1 astrocytes could be responding to direct stimulation by the environment immediately adjacent to plaques. Conversely, DAA2 astrocytes could reflect a chronic reactive state less dependent on ongoing stimulation by plaque-related pathological processes.

### Development of an integrated map of astrocytes from human single-nucleus data from AD, MS, and PD patients

To study the heterogeneity of human astrocytes in health and disease, we integrated astrocyte expression profiles from 13 human single-nucleus RNA-seq studies spanning three neurodegenerative diseases (AD, MS, PD) **(Table 2)**. Raw sequencing reads were obtained for each study and processed using the same pipeline as the mouse data, encompassing 995,000 astrocytes. We used a human reference dataset^52^ to annotate coarse cell types (see Methods). After rigorous QC, our integrated human atlas included a final total of 438,000 astrocytes. The high degree of dropout we observed resulted from removing clusters that scored high for markers of other brain cell types. This approach increases confidence that our dataset was highly enriched for astrocytes.

**Table 2.**
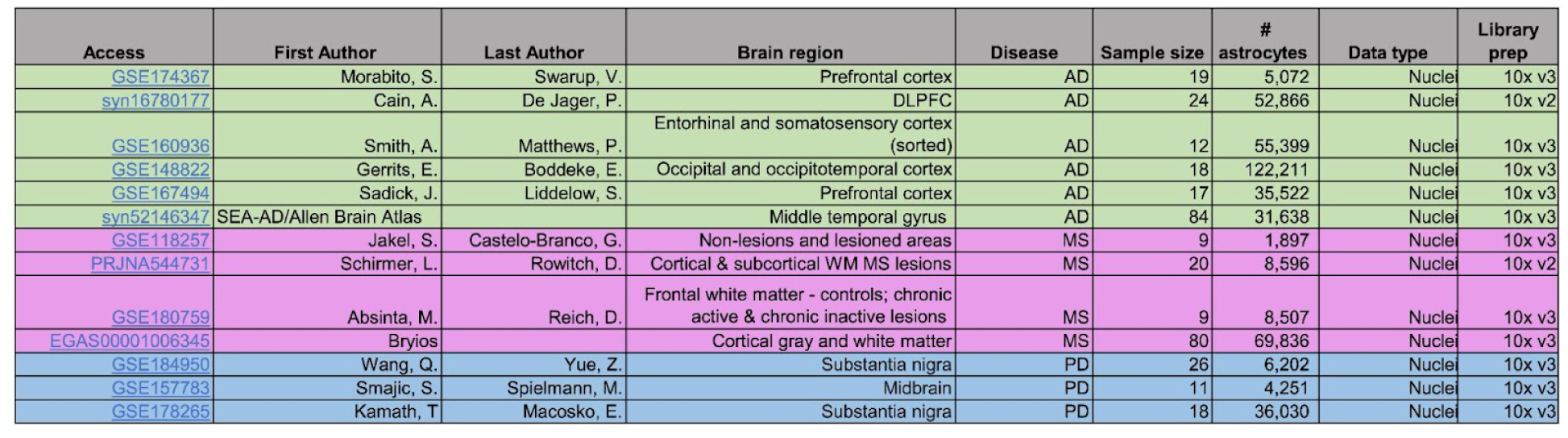
Table of datasets included in the human astrocyte atlas.

The AD studies included Prefrontal Cortex (60 donors), Entorhinal Cortex (12 donors), Occipital Cortex (18 donors), and Middle Temporal Gyrus (84 donors). The MS studies included cells from lesioned and non-lesioned areas of the nervous system (29 donors) and Cortical gray-white matter (80 donors). Lastly, the PD studies came primarily from Midbrain (11 donors) and Substania Nigra (37 donors). Metadata for each study, including the number of donors, brain region, and cell numbers, can be found in **Table 2**.

Using Harmony (see Methods), we generated an integrated space of human astrocytes (**Figure 5A**)^53–66^. Astrocytes were clustered using an iterative approach similar to that described in the mouse section above. Final clustering resulted in 8 distinct clusters with minimal study-driven batch effects, as evidenced by the presence of cells from each study within each cluster, and the contribution of samples from each study to each cluster (**Figure 5A-C, Supplemental Figure 4**) (Methods).

**Figure 5:**
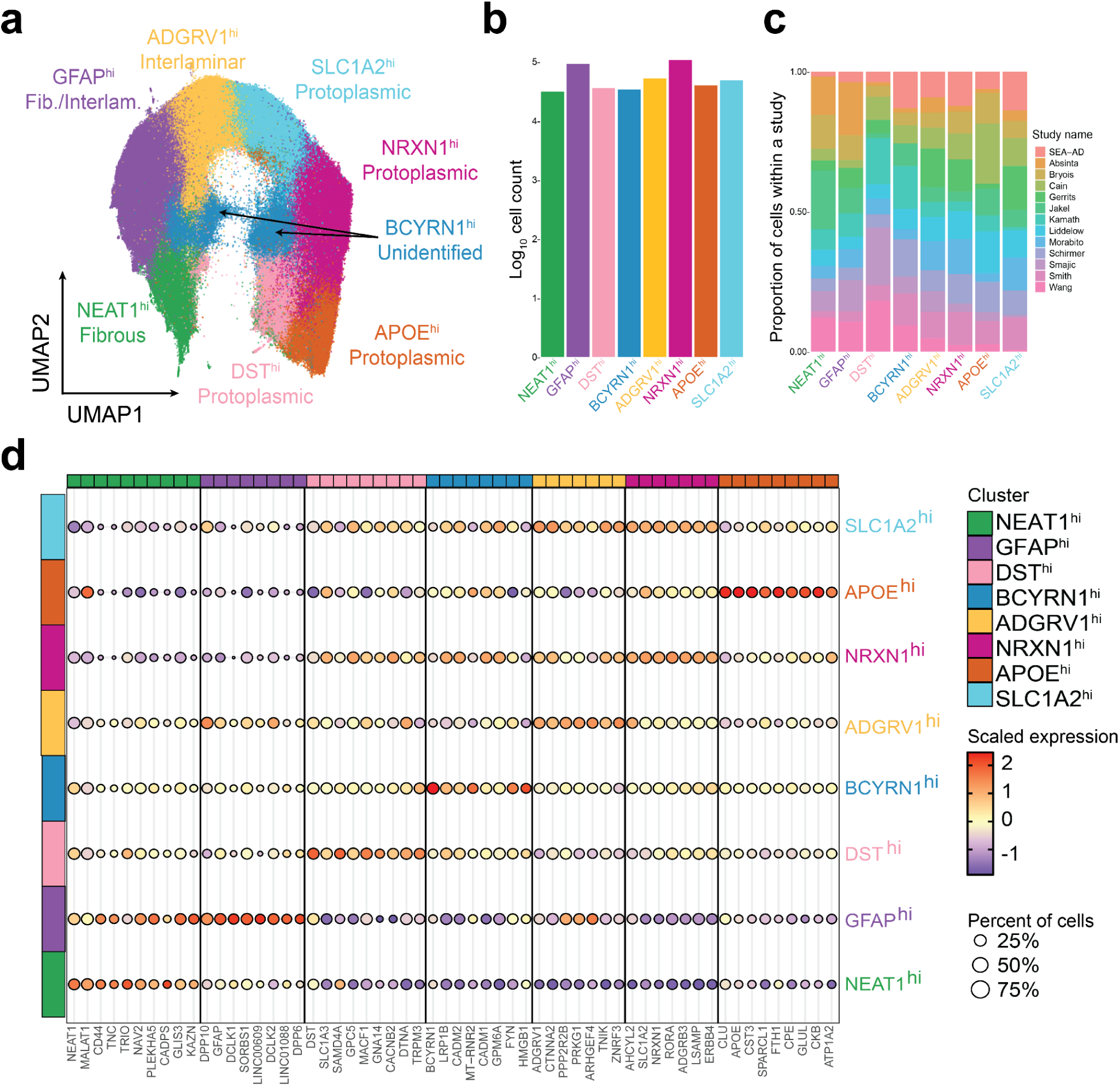
**Cell-level clustering of human astrocytes identifies broad astrocyte subtypes.** (A) Uniform Manifold Approximation and Projection (UMAP) of human neurodegeneration and control astrocytes revealing eight distinct clusters. (B) Counts of cells by cluster. (C) Proportion of cells per cluster, colored by study. (D) Dotplot of the top 10 cluster markers of each cluster. Color represents the scaled expression of each cell in that cluster. Dot size represents the percentage of cells that express that specific marker.

### Diverse astrocyte populations are detected in the integrated space

Human astrocytes have been described as divergent from their rodent counterparts, especially in specialized functions and morphology. Recent studies have shown that despite the general conservation of cellular architecture, there are extensive differences between homologous human and mouse cell types, including alterations in proportions, laminar distributions, gene expression, and morphology^67^. These species-specific features emphasize the importance of directly studying human brain samples.

Our human integrative analyses uncovered a broader diversity of astrocyte subtypes and populations than our mouse integration. In agreement with previous studies, based on marker gene expression we detected presumptive interlaminar astrocytes (with elevated ID3 and DPP10 ), protoplasmic astrocytes (lacking TNC, GFAP, and DPP10 with elevated SLC1A2/3), and fibrous astrocytes (expressing high GFAP and TNC) (**Figure 5A, 5D, Supplemental Figure 4A&B**) ^67^. In our atlas, we detected eight clusters, which we named for distinguishing marker gene expression. This included a fibrous astrocyte cluster that expressed *NEAT1*, which is linked to heightened astrocyte reactivity ^68^ (NEAT1-hi fibrous). Additionally, a separate GFAP-hi cluster, a well-characterized marker of astrocyte identity and activation, presented increased expression of ID3 and DPP10, suggesting a blend of fibrous and interlaminar characteristics (GFAP-hi Fib/Interlam). We also detected a population of interlaminar astrocytes with high levels of *ADGRV1* (ADGRV1-hi Interlaminar). Four populations of protoplasmic astrocytes were also detected. This included SLC1A2-hi and NRXN1-hi clusters, which also expressed high levels of *RORA.* RORA has been shown to have a neurosupportive role in astrocytes, directly transactivating the IL-6 gene. This direct control is necessary to maintain basal IL-6 levels in the brain^69^. We also identified an APOE-hi cluster of protoplasmic astrocytes that expressed high levels of *APOE* and *CLU*, well-established AD risk genes. Interestingly, a recent study showed that the CLU AD risk allele leads to increased CLU expression and enhanced inflammatory signaling in iPSC-derived astrocytes^70^. The fourth protoplasmic cluster we identified, DST-hi, expressed high levels of *DST* and *SAMD4A*, two genes shown to play a role in reactive astrocytes^71–73^. Lastly, we discovered a primate-specific BCYRN1-expressing cluster, reflecting the absence of this cluster in mice. While the BCYRN1 gene has been studied heavily in cancer, very little is known of its role in astrocytes. Combined, our integrative analysis identified transcriptionally diverse astrocyte populations, highlighting their varying functions in healthy and diseased brains.

### Astrocytes display disease-associated activation signatures

To characterize global gene expression changes in astrocytes in response to AD, MS, or PD pathology, we performed differential expression analyses using a similar meta-analysis approach as used for the mouse model analysis above (see Methods). We identified 577, 297, and 356 DEGs in AD, MS, and PD, respectively (**Figure 6 A-C**). The upregulated genes were, on average, largely replicated across datasets for each respective disease, pointing to the reproducibility of the activation signature across studies (**Figure 6 A-C**).

**Figure 6:**
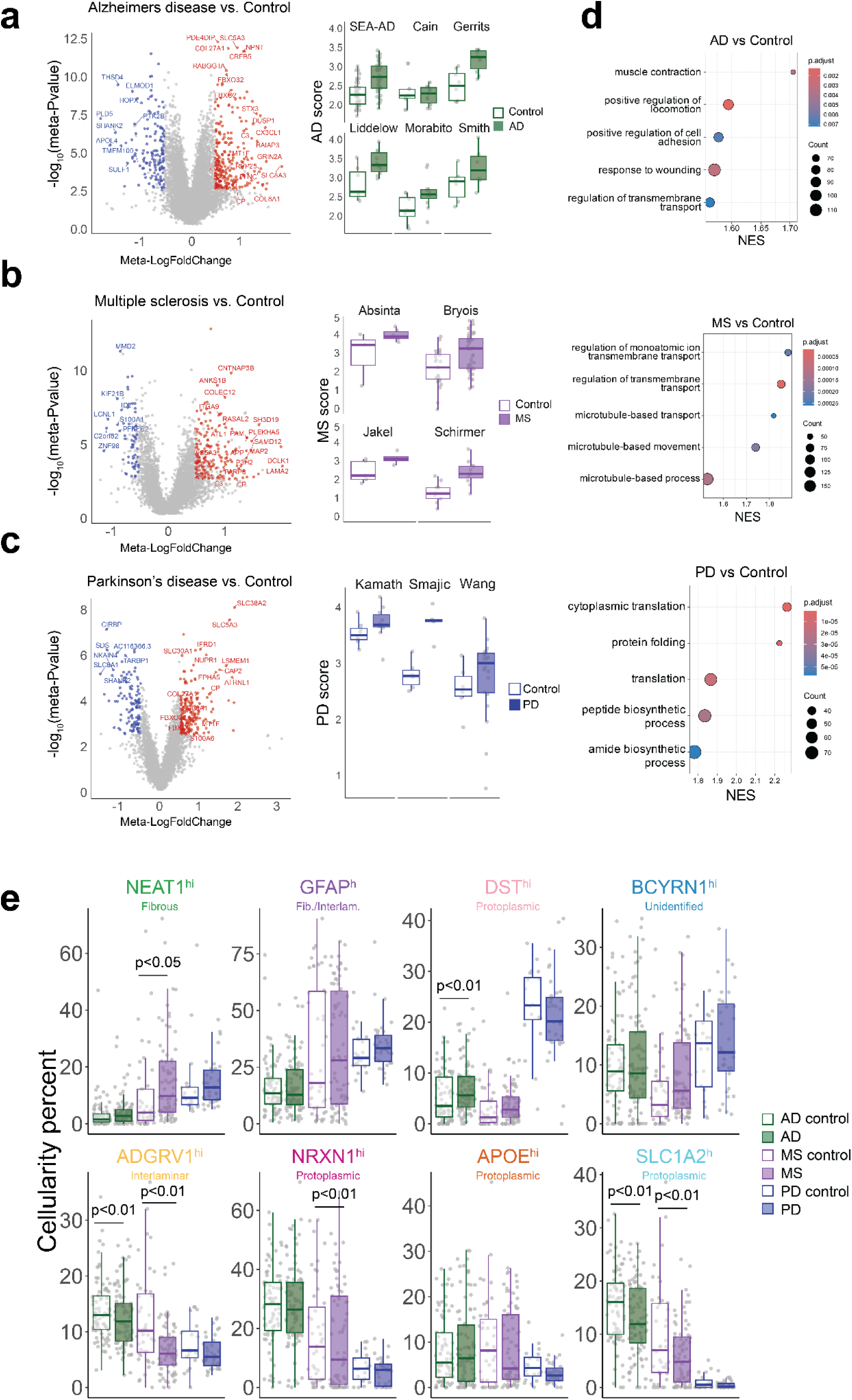
**Gene expression changes in human astrocytes across diseases** Differential expression analysis between disease and normal astrocytes for Alzheimer’s disease (A), Multiple Sclerosis (B), and Parkinson’s Disease (C). (left) Volcano plots for each differential expression analysis. Red and blue colors indicate significantly differentially expressed genes (meta-logfoldChange > 0.5, metaFDR < 0.05), and gray indicates no significant change (See Methods). (right) Boxplots showing average expression of upregulated Genes shown in the volcano plot, faceted by study. Dots represent an individual sample from the respective study and the y-axis represents the average expression of upregulated genes by disease. Study names are indicated at the top of each plot (see table 2 for details). (D) Gene Set Enrichment Analysis of Gene Ontology Biological Pathways. The color of the circles indicates the adjusted p-value, and the size indicates the Normalized Enrichment score for the specific pathway. Row names include the top scoring gene ontology categories. (E) Barplots indicating cellularity percent by study, cluster, and disease indication. Each dot represents a sample from an individual study. The y-axis represents the sample’s percentage of cells contributing to a specific cluster. Cluster names are indicated at the top of each plot. P-values are indicated over groups with significant differences in cellularity and were generated from the Kruskal-Wallis test and post-hoc Wilcox test on center-log transformed proportions(See Methods).

Transcriptional changes between cases and controls across each disease were largely distinct, with only 29 genes upregulated in all three diseases (**Supplemental Figure 4**). One example of such an upregulated gene is *CP,* which is largely produced by astrocytes in the brain and affects learning and memory in mice^74^. Another example, *SLC5A3,* was recently reported to be upregulated in mouse astrocytes following stroke^75^, potentially pointing to a basic activation profile for a subset of astrocytes reacting to diverse brain insults.

The transcriptional changes observed in AD were more closely correlated with those observed in PD (as opposed to MS; r=0.31 vs.0.02, respectively) (**Supplemental Figure 4D**). Additionally, genes consistently upregulated across AD and PD included collagens (*COL27A1* and *COL8A1*), proteins within the ubiquitination pathways (*FBXO2* and *FBXO32*), and metallothioneins (*MT1F* and *MT1G*) as well as *S100A6*, *SPARC*, *SLC38A2,* and others (**Supplemental Figure 4D**). The shared transcriptional changes between AD and MS were less numerous; however, some genes, like *C3,* were upregulated in both diseases^76^. PD and MS also shared some transcriptional changes not seen in AD, including upregulation of *SLC9B2* and ribosomal genes (*RPL32* and *RPL39*).

Although the gene-by-gene overlaps between diseases were minimal, a more sensitive approach using GSEA found that similar transcriptional programs were shared across diseases. For example, GSEA analysis of AD-associated genes identified an upregulation of pathways related to *regulation of transmembrane ion transport* and *response to wounding* (**Figure 6D, Supplemental Table 3**); MS-associated genes were also enriched in ion transport pathways (**Figure 6D**); PD-associated pathways were strongly enriched in protein folding and translation pathways, amongst others (**Figure 6D, Supplemental Table 3**). A comparative analysis of the enriched pathways highlighted some shared pathways between diseases. For example, AD and MS-associated genes were enriched in *regulation of transmembrane transport* (**Figure 6D, Supplemental Table 3**).

Next, we performed cluster abundance analyses to identify astrocyte sub-populations over- or under-represented across disease conditions (**Figure 6E**). We identified three clusters that were differentially abundant in AD, two that were depleted in AD, and one that was enriched. The two clusters depleted in AD samples were ADGRV-hi interlaminar astrocytes (p < 0.01) and SLC1A2-hi protoplasmic astrocytes (p = 0.003), suggesting these clusters might represent homeostatic astrocytes. The depletion of the SLC1A2-hi cluster was detectable across most studies, while the depletion of the ADGRV-hi cluster was driven by a subset of studies (SEA-AD, Cain, and Smith) **(Supplemental Figure 4A).** The single enriched cluster we identified was the DST-hi protoplasmic astrocytes (p < 0.01), suggesting these might represent a reactive AD-associated cluster. The expansion here was also driven by just a subset of studies (SEA-AD and Morabito) **(Supplemental Figure 4A)**.

Similar to AD, in MS, ADGRV-hi interlaminar astrocytes and SLC1A2-hi protoplasmic astrocytes were also significantly depleted (p < 0.001, p <0.001, respectively), supporting the interpretation of these clusters as homeostatic astrocytes. In addition, the NRXN1-hi cluster was also significantly depleted (p = 0.05) in MS. The depletion of SLC1A2-hi and NRXN1-hi clusters was relatively consistent across MS studies, while the depletion of ADGRV-hi was driven by a subset of MS studies (Bryois and Jakel, **Figure 6E**). At the same time, Neat1-hi fibrous-like astrocytes were expanded in MS (p = 0.02), suggesting that this cluster might represent an MS-reactive astrocyte population.

We did not identify any clusters significantly differentially abundant in PD, although we did note that expansion of NEAT-1 was trending towards significance (p = 0.07) (**Figure 6E)**. The lack of significant changes in abundance in PD samples could be related to lower power resulting from fewer studies and samples than for AD or MS Taken together, we identified robust pseudobulk transcriptional changes in astrocytes associated with neurodegeneration in AD, MS, or PD that were replicated across studies. Cluster abundance analysis highlighted putative disease-associated (NEAT1-hi and DST-hi) and homeostatic (ADGRV1-hi, NRXN1-hi, and SLC1A2-hi) astrocyte populations and substantial variability across studies.

### Expression of AD-associated genes is enhanced within the GFAP-hi astrocyte population

Given the complex changes occurring at both pseudobulk and cluster abundance levels, we wanted to investigate whether disease-specific DEGs were preferentially expressed in any of the astrocyte subtypes we identified. Interestingly, AD DEGs showed the most distinct expression pattern and were enriched in GFAP-hi astrocytes (**Figure 7A-B; Supplemental Figure 4C).** To further investigate this population, we subclustered these GFAP-hi astrocytes, which resulted in four additional subpopulations (**Figure 7C**). We discovered both fibrous-like (*NEAT1*^hi^/*TNC*^hi^ SubCluster 2) and interlaminar (*DPP10*^hi^ SubCluster 4) within the *GFAP-hi* cluster as well as two additional subpopulations (*CTNND2*^hi^ astrocytes, SubCluster 3 and *HSP90AA1*^hi^ astrocytes, SubCluster 1) (**Figure 7C, Supplemental Figure 5A**). Interestingly, Subcluster 1 was trending towards expansion in AD (p = 0.054), with most studies showing a moderate to considerable increase in this population in AD. While Subcluster1 was not differentially abundant between disease and normal states, the transcriptional program in this subset of GFAP-hi astrocytes may reflect disease-specific activation states. Also of note, subcluster 4 (DPP10-hi) exhibited significant enrichment for pathways related to cilia assembly and organization **(Supplemental Figure 5B)**. This subcluster also had elevated expression of FOXJ1 and SPAG17, both key markers associated with ciliary structure and function. These findings align with a recent study identifying SPAG17+ ciliated astrocytes in MS lesion contexts^43^. Within our study, the prominence of these markers in subcluster 4 suggests a specialized function for these astrocytes in cilia-associated pathways, potentially contributing to cellular signaling and environmental modulation in neurological conditions.

**Figure 7.**
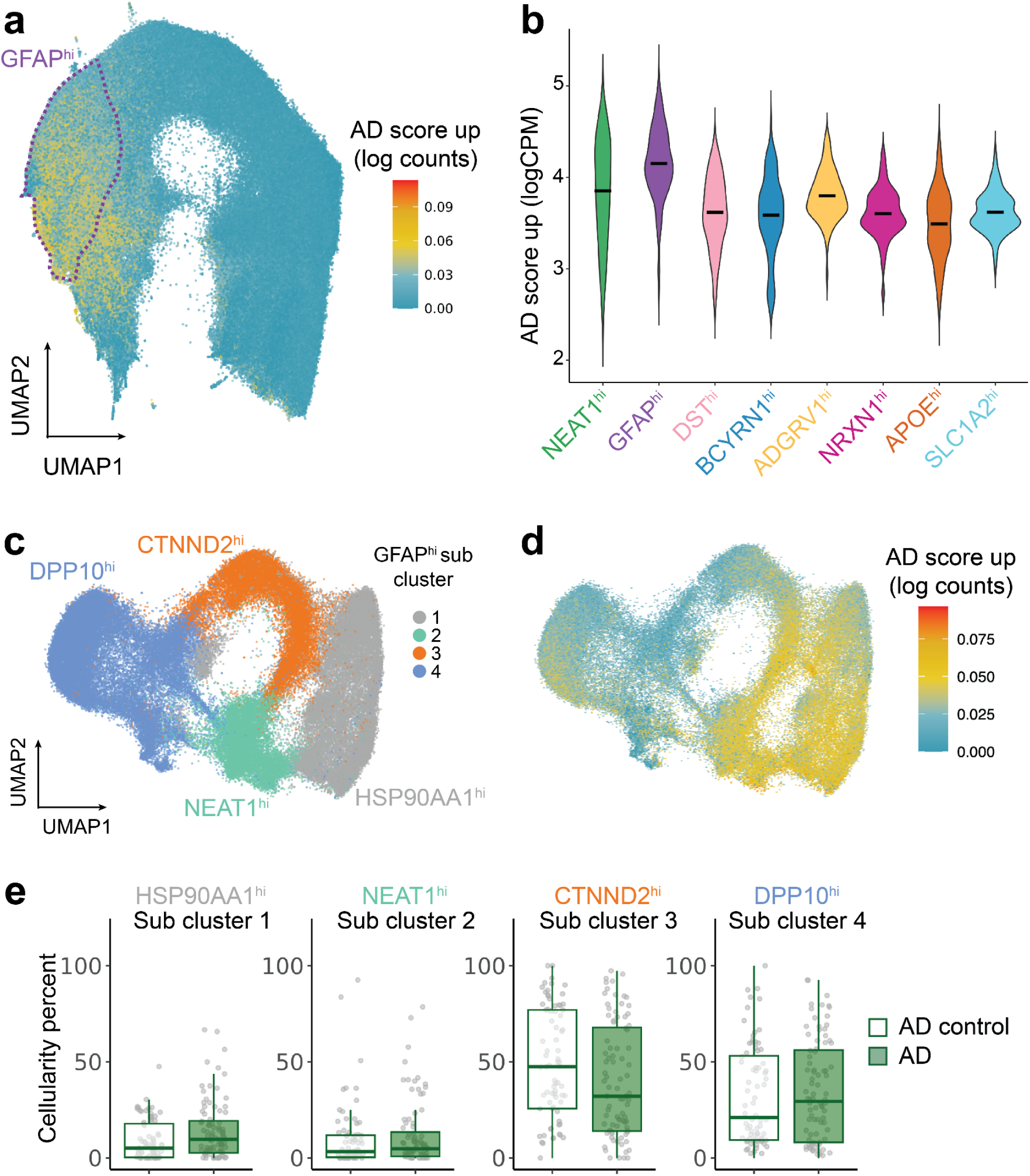
**AD-associated genes are enriched in GFAP-hi Astrocytes.** *(A)* Scoring human integrated neurodegeneration astrocytes with differentially expressed genes between AD and control astrocytes (logFC>0.5 p <0.05). UMAP indicates the distribution of expression scores. Dotted line indicates the GFAP-hi cluster. *(B)* Violin plot of AD score, by cluster. (C) UMAP showing subclustering of the human GFAP-hi cluster. Iterative clustering resulted in four distinct subclusters. (D) Human GFAP-hi subclusters scored by differentially expressed genes from AD astrocytes. (E) Barplots indicating cellularity percent by cluster and disease indication. Each dot represents a sample from an individual study. The y-axis represents the sample’s percentage of cells contributing to a specific cluster. Cluster names are indicated at the top of each plot. P-values are indicated over groups with significant differences in cellularity and were generated from the Kruskal-Wallis test and post-hoc Wilcox test on center-log transformed proportions.

### Transcriptional programs induced in human disease are partly recapitulated by mouse models of disease

To determine how our clustering and integration in mice and humans compare, we first wanted to compare global differential gene expression changes across species and by disease. To this end, we examined the concordance of gene expression changes in human disease and mouse models of AD **(Figure 8A & B)** and MS **(Figure 8C & D)**. The comparison of significantly upregulated astrocyte genes in AD or MS mouse models vs patients revealed a large distinction between the species, with only 35 genes in common in AD and 4 in MS. Interestingly, one of the genes most strongly induced in both mouse and human AD samples is *C3*, a central component of the complement signaling pathway that mediates synapse loss and neurodegeneration in mouse models of AD ^77,78^. Furthermore, we observed a shared upregulation of other genes likely to be involved in microglia communication and astrocyte activation (*CX3CL1*, *C1QL1*)^79–81^, neuroprotection and repair (*CP*, *S1PR3*)^82,83^, and genes linked to astrocyte-induced inflammation (*ITGA5*, *IGFBP5*)^84–86^. In MS, although fewer genes had consistent differential expression patterns between species, we identified differential expression of canonical reactive astrocyte genes related to a pro-inflammatory state (*TMSB4X*, *VIM*)^1,49^. The fact that we identified fewer similarities between mouse models of MS and human disease may suggest that mouse models of MS included in this atlas incompletely model the full spectrum of human astrocyte disease responses.

**Figure 8.**
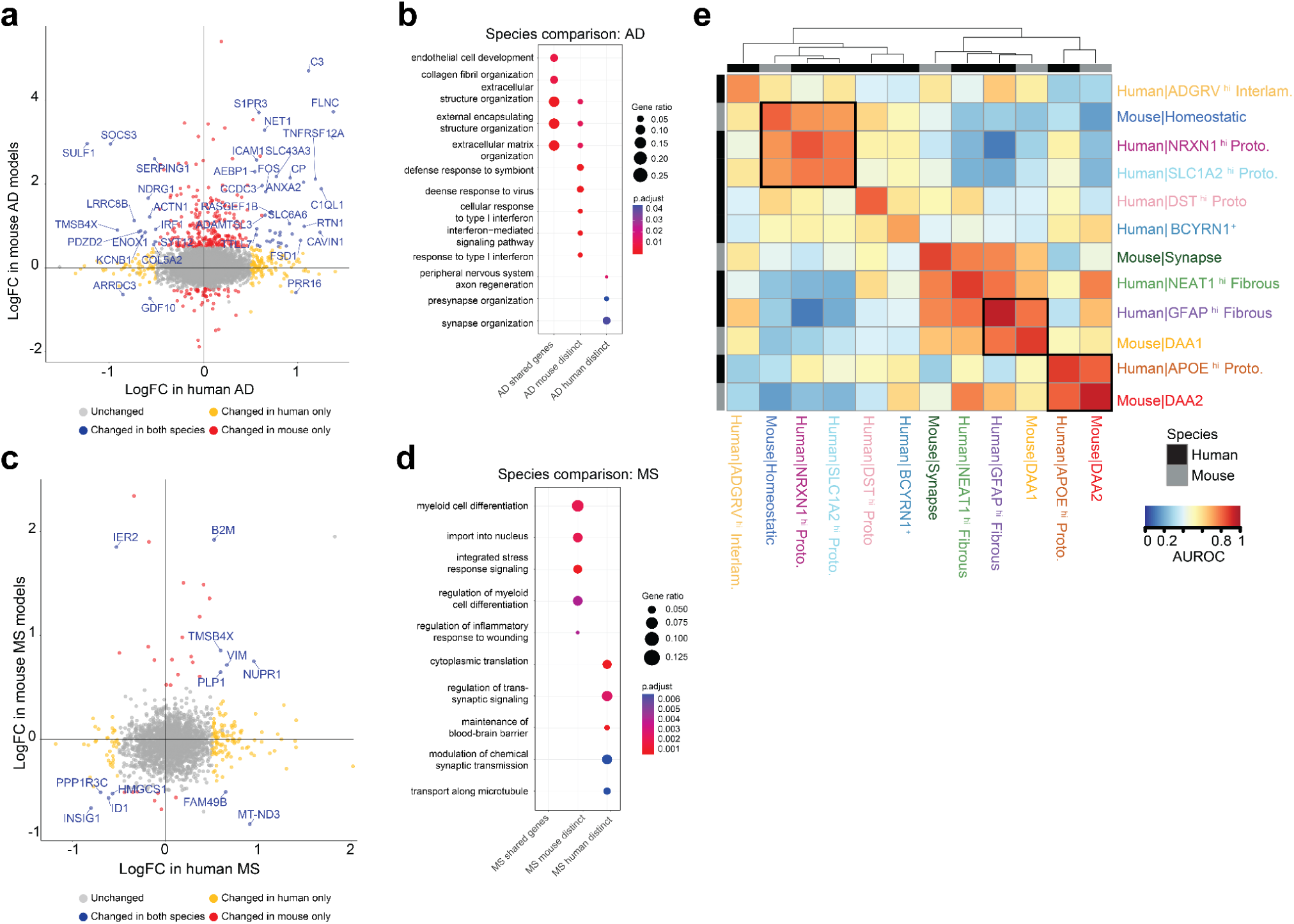
**Cross-species comparison reveals species-specific gene expression changes in disease but conservation of subpopulations.** Comparison of differentially expressed genes between human and mouse astrocytes AD vs. control (A), and MS vs. control (C). The y-axis represents log-fold changes in mouse disease models vs controls, and the x-axis represents log-fold changes in human patients vs controls. Color represents significantly changed genes in both species (blue), only in humans (red), only in mice (yellow), or not changed in either species (gray). Dotplot of Gene Ontology overrepresentation analysis of shared and distinct differentially expressed genes between mouse and human in AD (B) and MS (D). The color of the circles indicates the adjusted p-value, and the size indicates the ratio of genes in that pathway. (E) Heatmap of the mean area under the receiver operator characteristic (AUROC) from analysis of human and mouse clusters by MetaNeighbor. Colors represent the strength of the comparison, with red values reflecting more similarity between cells, and blue values reflecting less similarity between cells. Top and side annotations represent clusters from human (black) or mouse (gray). Cluster names are labeled on both columns and rows. Black boxes indicate similar clusters across species that are highlighted in the text.

To further characterize the distinct and shared responses across species, we conducted GO enrichment analyses on the shared and distinct genes in AD and MS **(Figures 8B & D)**. In AD, the shared genes pointed primarily to extracellular organization, highlighting the likely morphological response commonly seen in astrocytes in both humans and mice in response to neuropathologies. A notable species-specific divergence was observed; mouse astrocytes showed an interferon response, while human astrocytes predominantly upregulated genes associated with synaptic organization **(Figure 8B)**. In the case of MS, the lack of sufficient shared upregulated genes precluded a robust enrichment analysis. Nevertheless, the distinct pathways revealed similarities to AD; mouse astrocytes showed a pronounced immune and inflammatory response, while human astrocytes upregulated synapse signaling-related genes **(Figure 8D)**. These findings highlight the critical role of astrocytes in neuroinflammation and synaptic regulation and suggest that these cells may play different roles in human disease vs mouse models. At the same time, within species, global gene expression changes seen in astrocytes across diseases are actually highly conserved.

### Cross-species comparison of astrocyte clusters reveals correlated subpopulations

To evaluate the conservation of astrocyte clusters between mice and humans, we leveraged MetaNeighbor to assess cell-type replicability across our integrated dataset ^87^. This approach has been used previously for inter-species comparisons^88^. We quantified transcriptional similarities by using the mean area under the receiver operator characteristic (AUROC) scores, enabling us to use hierarchical clustering to sort clusters with similar AUROC scores **(Figure 8E)**. High AUROC values between clusters demonstrate striking cross-species similarities.

Notably, the mouse Homeostatic astrocytes cluster closely with human protoplasmic astrocyte clusters, NRXN1-hi and SLC1A2-hi. All three of these clusters are significantly reduced in disease samples, which would be consistent with all three functioning in homeostatic roles.

Further, the mouse DAA populations cluster with distinct human populations. The DAA1 population clusters most closely with human fibrous GFAP-hi astrocytes. Across these human and mouse clusters, we noted enrichment not only of GFAP, but of other canonical reactive astrocyte genes such as VIM and ID3. On the other hand, the mouse DAA2 population clusters most closely with human protoplasmic APOE-hi astrocytes. In both human and mouse samples, these astrocytes are distinguished by the expression of AD-associated genes, APOE and CLU, which could point to a causal role in disease biology. Together, these data highlight that the foundational astrocyte states are conserved across species and emphasize the shared biological frameworks that persist amidst species-specific adaptations **(Figure 8E)**.

## Discussion

Here, we present a comprehensive meta-analysis of astrocytes across species and diseases, leveraging the massive amount of publicly available single-cell datasets. Our analysis of mouse samples identified clusters of astrocytes, including Homeostatic, Synapse-related, and two distinct disease-associated populations, DAA1 and DAA2. A detailed characterization of DAA1 and DAA2 highlighted transcriptional differences between these populations as well as differences in their spatial proximity to disease pathology. These data point to potentially divergent biological functions. Specifically, DAA1 astrocytes are proximal to amyloid plaques and have an expression profile resembling acute activation by LPS, while DAA2 are distal to amyloid plaques and have an expression profile that is more distinct from acute activation.

Further understanding the differences between these two subtypes may offer insights into the spatiotemporal dynamics of astrocytes in plaque-rich vs. plaque-distal regions, which could be crucial to understanding AD progression and inspire therapeutic strategies targeted toward beneficial versus harmful astrocyte responses.

In parallel to the characterization of mouse astrocytes, we performed an analysis of human tissue samples, capturing the three categorical subtypes of astrocytes: protoplasmic astrocytes, which populate the grey matter and are the most abundant subtype; fibrous astrocytes; and interlaminar astrocytes. Interestingly, while human astrocytes do not exhibit universal disease programs as seen in mouse samples (**Supplemental Figure 4D**), subtype-specific disease-enriched populations were identified (**Figure 6E**). Most gene expression changes occurring in disease were observed in the fibrous/interlaminar GFAP-hi subtype of astrocytes, particularly in Alzheimer’s Disease (AD) and Parkinson’s Disease (PD) **(Figure 7A, Supplementary** Figure 4C), suggesting that these astrocytes may be more reactive or vulnerable to pathological stimuli than other subtypes. This is in contrast with our analysis of MS samples, where gene expression changes were observed across multiple astrocyte subtypes.

Of particular note is the high correlation between the mouse DAA1 and DAA2 populations, and the human fibrous and a subset of protoplasmic astrocytes, respectively, pointing to shared biological processes across these cell types. Intriguingly, the DAA2 cluster and corresponding protoplasmic cluster both express high levels of APOE and CLU, two well-known AD GWAS-associated disease loci. This could suggest that these populations of cells play key roles in the pathogenesis of AD. Our analysis comparing integrated human and mouse astrocyte subclusters highlight the power of our atlas to aid in identifying and characterizing preclinical models most relevant to studies of human disease. At the same time there was overall low correlation between disease DEGs in mouse vs human astrocytes, especially in MS, suggesting caution in interpreting the detailed roles of astrocytes in disease based solely on preclinical models. However, the partial overlap observed in AD models and patient AD samples indicates that certain aspects of mouse astrocyte function could be relevant for understanding astrocyte biology in human samples.

Our analysis highlights a number of insights into astrocyte subcluster biology that are concordant with other recent data sets. For example, in our dataset we observed that mouse Homeostatic and Synapse-related and human (SLC1A2-hi and NRXN1-hi protoplasmic) homeostatic clusters are the highest expressers of the synaptic neuron and astrocyte program (SNAP) **(Supplemental Figure 5C&D)**. The SNAP program includes astrocyte genes involved in synaptic functions and was observed to decline both by age and also in patients with schizophrenia^89^ . The identification of SNAP expression in these cells is consistent with SNAP expression being beneficial, and the loss of these cells in neurodegeneration likely leading to disrupted astrocytic support of neurons. Another interesting observation from our analysis is that our NRXN1-hi protoplasmic cluster expressed similar marker genes and proportional changes as those in the Astro-2 population identified in the Seattle Alzheimer’s Disease Brain Cell Atlas^90^. This points to the high reproducibility of a particular subcluster of protoplasmic astrocytes, that is specifically lost during disease, expresses SLC1A2 and NRXN1, and performs homeostatic functions in the brain.

A recent atlas of cellular communities in the prefrontal cortex during aging and in AD identified an astrocyte subpopulation, Ast.10, implicated by causal modeling in mediating the effects of Tau on cognitive decline ^91^. Our analysis highlighted that the Ast.10 cluster from this study has transcriptional similarities to both DAA1 and DAA2, including expression of Mt1 and Mt2 **(Supplemental Figure 5C&D)**, supporting that further analysis of the programs in these cells could reveal novel disease-relevant functions. In humans, we see increased expression in the APOE-hi cluster. Another recent publication mapped astrocyte transcriptomic changes along the spatiotemporal progression of AD^92^. They identified two reactive populations, astR1 and astR2 that increased in proportion during disease progression. Similar to our NEAT1-hi and GFAP-hi fibrous clusters, astR1 and astR2 marker genes included *GFAP*, *AQP4, CD44* and *TNC*. Additionally, a subset of GFAP-hi astrocytes also shared expression of stress response genes like *HSP90AA1.* The commonalities observed across distinct efforts to catalog aspects of astrocyte state in disease and our integrated analysis validates the utility and strength of our atlas in interrogating astrocyte biology.

Our study significantly enhances our understanding of astrocyte heterogeneity by integrating data from multiple neurodegenerative diseases, including AD, multiple sclerosis (MS), and Parkinson’s disease (PD). Through cross-species analysis, we highlight both conserved and disease-specific astrocyte clusters, showcasing the functional diversity of these cells across different pathological contexts. The distinct and disease-specific nature of astrocyte populations highlights the importance of considering cell-type-specific responses in neurological diseases. The annotation of cell types across species with high resolution advances the translational understanding of astrocytes in healthy and neurodegenerative contexts. Similarities between mouse and human DAAs underscores the utility and limitations of mouse models in studying human neurological diseases. Our atlas (http://research-pub.gene.com/AstroAtlas/) will be valuable for guiding future research using targeted experimentation to elucidate the specific roles of varying astrocyte subtypes in these disease contexts and developing appropriate therapeutic interventions.

## Methods

### Mouse data preprocessing and clustering

A total of 15 mouse datasets were analyzed spanning 6 AD and 3 MS models, and 181 samples. Mouse and human datasets included in this study are listed in Tables 1 and 2. We collected publicly available FASTQs from Gene Expression Omnibus (GEO) with GSE IDs, as indicated in Table 1. Additionally, we retrieved metadata for each sample, including mouse strain, age, genotype, and treatment, and technical variables, including library preparation, sequencing protocol, or any batch structure, when provided. FASTQs were processed through the cellranger pipeline if 10X data or the Drop-seq pipeline if drop-seq data ^93,94^, using default parameters in both cases. For 10X samples, ambient RNA was identified and filtered using *Cell Bender*. For Drop-seq data, ambient RNA was filtered using *SoupX* on unfiltered counts tables. Each dataset was processed separately using *Seurat* standard processing protocols. Briefly, we calculated the number of unique features (*nFeature*), total number of unique UMIs, percentage of mitochondrial genes (*percent*.*mito*), and percentage of ribosomal genes (*percent.ribo*). Additionally, doublets were identified using *scDoubletFinder*.

To identify astrocytes, we scored each cell for hallmark brain cell gene markers. The final cells included from each study were cells labeled as astrocytes through marker expression and met all other QC metrics (*nGene* > 200, *percent.mito* < 25%, *cell.class* == Singlet, *microglia/oligo/OPC/neuron/BMEC/VSMC score* < 0.5 ).To ensure consistent identification of astrocytes within each sample we used a label transfer of cell labels from GSE15389 using SingleR. Supplemental Table 1 summarizes the final dataset by study. For each dataset, data was normalized, and the top 2000 variable genes were identified using the VST selection method.

We performed integration using *Seurat Canonical Correlation Analysis,* using the 2000 most variable genes from each study as anchors. Using the integrated experiment object, we generated UMAPs and PCA using the 2000 anchor features.

Integration quality was assessed using multiple approaches. First, we generated a Uniform Manifold Approximation and Projection Map (UMAP) of the integrated space and conducted a visual assessment of the projection of cells by study. In addition, we conducted a quantitative analysis. Here, we divided the UMAP space into 12 bins. Within each bin, we assessed whether any study was over-represented in a specific bin using a hypergeometric test. We conducted this test split unbiasedly across a various number of bins as well as other sample information bins, including data types(single cell/single nuclei), library preparation (10x/drop-seq), and brain region (hippocampus/cortex/hemisphere). Data from this analysis can be found in supplementary figure 1.

We used Seurat’s FindClusters function to iterate over a range of clustering resolutions, from low (*res=0.05)* to high (*res=1*), to determine an appropriate *number* of clusters. At each resolution, cluster markers were generated using *scoreMarkers.chan* from the *scran.chan* package in R. The final resolution, resulting in 4 clusters, was used for the analysis as it was the lowest resolution and produced at least 20 significant (*Cohen.mean* > 0.5) markers per cluster.

### Human data preprocessing and clustering

We aggregated 16 human datasets from AD, MS, PD, and Control tissues. Each dataset was individually preprocessed. *perCellQCMetrics.chan()* and *perCellQCFilters.chan()* function from *scran.chan* package was used to filter out low-quality cells (either due to low UMI counts, low detected genes, or high percent mitochondrial UMI). Astrocyte-labeled cells were then extracted from each dataset. All astrocyte profiles were then merged into a single counts matrix and *multiBatchNorm(normalize.all = TRUE, batch = batch)* function from the *batchelor* package was used to compute batch-corrected normalized log-expression values. To avoid capturing batch effects, the top 2000 highly variable genes were identified in every dataset individually using *modelGeneVar.chan* function; highly variable genes were then ranked based on the number of datasets they were identified in and their median variance across datasets; consensus top 2K variable genes were then selected and used for downstream analysis. PCs were computed using fixedPCA function from scran package. Initial integration was performed using a *runHarmony(theta=2)* function from the *harmony* package. UMAP was computed using *runUMAP* function from the *scater* package followed by *clusterSNNGraph.chan* from *scran.chan* package with default parameters to obtain initial clustering of the integrated space. To remove potential non-astrocytic cells, we scored the space based on canonical markers of other brain-resident cell types. We identified small clusters that likely represented microglia and perivascular macrophages, ependymal cells, oligodendrocytes, and neurons. Donors with less than 20 cells were also removed. After removing contaminating clusters, variable genes, PCs, UMAP, and integrated space (*runHarmony(theta=1.5*)) were recomputed using the same approach described above. Clusters were computed at varying resolutions from 0.1 to 1 with an increment of 0.1 using the *clusterSNNGraph.chan* function. A resolution at which each cluster is significantly different from all others (cohen.mean > 0.5) by at least one marker. Resolution of 0.5 was selected in this manner.

### Cellularity and Differential Abundance Analysis

Cellularity results were obtained by computing cluster abundances per donor and using a *cpmByGroup* function from the *edgeR* package to obtain normalized counts per cluster per donor that were then visualized as percentages per cluster in Figures 2 and 4. To test if any of the clusters were associated with a disease state, we performed differential abundance testing. Since cellularity is compositional, we transformed the proportions using a Center log-ratio (CLR) transformation. We then performed a kruskal-wallis non-parametric significance test on each cluster. Significant clusters were identified as significant after p-values were adjusted for multiple hypothesis testing (pVal/ #clusters tested). Significant clusters were followed up with a post-hoc Wilcox test between disease and control samples. The resulting p-values were FDR corrected, and clusters that passed an FDR 5% cutoff were deemed significantly different.

### Differential Gene Expression Analysis

A challenge in determining meaningful changes in large meta-analyses is the heterogeneous nature of the data, which can lead to false positives and negatives due to the effect of outliers. Therefore, we performed differential expression through the random effects approach described in ^41^. This method assumes that the different studies are estimating different, yet related, effects due to disease conditions. Differential expression results are calculated on pseudo-bulked cells, by sample, for each study separately, using *edgeR*. The standard errors of the study-specific *logFC* estimates are adjusted to incorporate a measure of the extent of variation, or heterogeneity, among the disease effects observed in different studies using the *metafor* package in *R*. The amount of variation, and hence the adjustment, can be estimated from the disease effects and standard errors of the studies included in the meta-analysis. The significance of the effects reported using this random effects model was calculated using Fisher’s combined probability test similarly and carried out using the *metap* package in *R*. For both mouse and human studies, significant genes were identified as those genes where *|dl_mu|* >= 0.5, *FDR* <=0.05, with the directionality of the effect estimated as positive for upregulated genes or as negative for downregulated genes in at least half the studies for a given disease. Given the higher variability of the human data, we required that upregulated and downregulated genes had a positive or negative estimated effect in all of the studies in MS (4 studies) and PD (3 studies). Given that we had more AD studies included (7 studies), we required that 4 out of the 7 studies had an estimated effect size of |dl_mu| >= 0.5.

### Gene Set Enrichment Analysis

Differentially expressed genes by disease or by cluster were ordered by effect size (“beta” for DE analysis, LogFC for all others). Genesets from GO Biological Process, GO Molecular Function, and KEGG were retrieved from the *clusterProfiler* package in *R* ^37^. To identify enriched pathways for each database we conducted Gene Set Enrichment Analysis for the corresponding database (*gseGO*, *gseKEGG*) with default settings, limiting geneset sizes to a minimum of 10 genes.

### Mouse LPS Single-Cell Study

#### Animals

Male C57BL6 (Charles River Hollister) aged 4-5 months were injected intraperitoneally with PBS vehicle control or LPS (1mg/kg) n=5 for each group. All protocols involving animals were approved by Genentech’s Institutional Animal Care and Use Committee, following guidelines that adhere to and exceed state and national ethical regulations for animal care and use in research. All mice were maintained in a pathogen-free animal facility under standard animal room conditions (temperature 21 ± 1°C; humidity 55%–60%; 12h light/dark cycle).

### Mice Perfusion and preparation of single-cell suspensions

48 hours post-injection, mice were perfused with ice-cold PBS, and the hippocampi were immediately sub-dissected. Single-cell suspensions were prepared from the hippocampi as described by. Briefly, hippocampi were chopped into small pieces and dissociated with enzyme mixes in Neural Tissue Dissociation Kit (P) (Miltenyi 130-092-628) in the presence of actinomycin D. After dissociation, cells were resuspended in Hibernate A Low Fluorescence medium (Brainbits) containing 5% FBS, with Calcein Violet AM (Thermo Fisher C34858) and propidium iodide (Thermo Fisher P1304MP). Flow cytometry was used to sort and collect live single-cell suspensions for the single-cell RNA-seq study.

### Single-cell RNA-seq library preparation and sequencing

Sample processing and library preparation were carried out using the Chromium Next GEM Automated Single Cell 3’ Library & Gel Bead Kit v3.1 (10X Genomics) according to the manufacturer’s instructions. Cells were prepared to aim for 10,000 cells per sample, and libraries were sequenced with HiSeq 4000 (Illumina).

#### Analysis

FASTQ files were analyzed with an in-house pipeline incorporating Cell Ranger to count and CellBender to filter ambient RNA. Sample quality was further assessed based on the distribution of per-cell statistics, such as total number of reads, percentage of reads mapping uniquely to the reference genome, percentage of mapped reads overlapping exons, number of detected transcripts (UMIs), number of detected genes, and percentage of mitochondrial transcripts using Seurat. Finally, astrocytes were identified as described above and filtered for comparative analysis.

Differential Expression analysis was performed using psuedobulk astrocytes by sample and run through a standard voom-limma workflow to generate log fold-changes and p values. Significant genes were labeled as those with absolute value logFC > 1 and p < 0.05.

#### Spatial Profiling of Astrocytes using RNAscope

The Institutional Animal Care and Use Committee (IACUC) at Genentech approved all experimental procedures involving transgenic animals. Tissue harvest, in situ hybridization, and image analysis were performed as described in Rao. et. al. STAR Protocols^50^. Briefly, 10-12-month-old wild type and TauPS2APP were anesthetized with an intraperitoneal injection of 2.5% Avertin. At least 3 mice of each genotype were used for experimental analysis. Animals were transcardially perfused with ice-cold phosphate-buffered saline and hemibrains were dissected, immediately embedded in cryoprotectant, and stored at -80℃. Coronal 5-7µm tissue sections were collected to perform in situ hybridization using commercially available RNAscope™ probes followed by immunohistochemistry for Aβ plaques (anti-β-amyloid 1-16 6E10, mouse, Biolegend cat# 803003). Aβ signal was amplified using a horseradish peroxidase secondary antibody, followed by TSA-Digoxigenin labeling and detection with an anti-DIG antibody (Opal 780 Reagent Pack, Akoya Biosciences, cat# FP1501001KT). Sections were mounted with Prolong Gold (Millipore Sigma, Cat#P36930) and imaged within a week. All samples were imaged using an Olympus VS200 slide scanner equipped with an X-Cite XYLIS XT720S LED light source (Excelitas). Wide-field fluorescent Z-stacks were acquired using a 20x air-objective, with a 5µm axial range, 0.5µm, and 0.325µm axial and X-Y resolution, respectively. DAPI, FITC, TRITC, Cy5, and Cy7 filter sets (Excelitas) were used to capture 16-bit fluorescent images. Single fluorophore, and fluorophore-minus-one controls were generated for each fluorophore used to determine optimal exposures and absence of spectral bleed-through. Images of coronal sections were processed by segmenting brain sections by anatomical region (hippocampus, cortex, and white matter) using the Allen Brain Atlas as a reference. Cell boundaries were approximated with 5µm expansion around DAPI-positive nuclei and mRNA puncta and plaque signal by respective fluorophore signal using QuPath (https://qupath.github.io/). Data was exported and analyzed in R Studio (http://www.rstudio.com/) to calculate mRNA puncta detected per cell for all samples.

Nearest-neighbor analysis was performed to determine how cell types changed expression of genes across tissue sections. Once cells were identified and gene expression was assessed in each cell, expression was normalized to cell size by dividing total expression by cell area.

Normalized expression was used to determine if a cell was indeed an astrocyte and which subtype was used. Various expression cutoffs were used, and final expression cutoffs were determined based on assumed proportions of astrocytes in tissue using reference data, while subtype cutoffs were based on assumed proportions from our integrated map. Each cell in transgenic animals was then assessed in terms of its distance to the nearest plaque using the physical (euclidean) distance from the cell center to the plaque center. Lastly, the total number of astrocytes was then totaled at each binned distance away from plaques, and subtype ratios were calculated and plotted.

## Supporting information

All Supplemental Figures

## References

1. Zamanian, J. L. et al. Genomic Analysis of Reactive Astrogliosis. J. Neurosci. 32, 6391–6410 (2012).

2. Cekanaviciute, E. & Buckwalter, M. S. Astrocytes: Integrative Regulators of Neuroinflammation in Stroke and Other Neurological Diseases. Neurotherapeutics 13, 685–701 (2016).

3. Liddelow, S. A. et al. Neurotoxic reactive astrocytes are induced by activated microglia. Nature 541, 481–487 (2017).

4. Brosnan, C. F. & Raine, C. S. The astrocyte in multiple sclerosis revisited. Glia 61, 453–465 (2013).

5. Wyss-Coray, T. et al. Adult mouse astrocytes degrade amyloid-β in vitro and in situ. Nat. Med. 9, 453–457 (2003).

6. Rossi, D. & Volterra, A. Astrocytic dysfunction: Insights on the role in neurodegeneration. Brain Res. Bull. 80, 224–232 (2009).

7. Liddelow, S. A. & Barres, B. A. Reactive Astrocytes: Production, Function, and Therapeutic Potential. Immunity 46, 957–967 (2017).

8. Eng, L. F., Vanderhaeghen, J. J., Bignami, A. & Gerstl, B. An acidic protein isolated from fibrous astrocytes. Brain Res. 28, 351–354 (1971).

9. Escartin, C. et al. Reactive astrocyte nomenclature, definitions, and future directions. Nat. Neurosci. 24, 312–325 (2021).

10. Sofroniew, M. V. Molecular dissection of reactive astrogliosis and glial scar formation. Trends Neurosci. 32, 638–647 (2009).

11. Guttenplan, K. A. et al. Neurotoxic reactive astrocytes induce cell death via saturated lipids. Nature 599, 102–107 (2021).

12. Jiwaji, Z. & Hardingham, G. E. Good, bad, and neglectful: Astrocyte changes in neurodegenerative disease. Free Radic. Biol. Med. 182, 93–99 (2022).

13. Habib, N. et al. Disease-associated astrocytes in Alzheimer’s disease and aging. Nat. Neurosci. 23, 701–706 (2020).

14. Jiwaji, Z. et al. Reactive astrocytes acquire neuroprotective as well as deleterious signatures in response to Tau and Aß pathology. Nat. Commun. 13, 135 (2022).

15. Al-Dalahmah, O. et al. Single-nucleus RNA-seq identifies Huntington disease astrocyte states. Acta Neuropathol. Commun. 8, 1**9** (2020).

16. Halpern, M., Brennand, K. J. & Gregory, J. Examining the relationship between astrocyte dysfunction and neurodegeneration in ALS using hiPSCs. Neurobiol. Dis. 132, **10**4562 (2019).

17. Kushwaha, R., Sinha, A., Makarava, N., Molesworth, K. & Baskakov, I. V. Non-cell autonomous astrocyte-mediated neuronal toxicity in prion diseases. Acta Neuropathol. Commun. 9, 22 (2021).

18. Lee, H.-G. et al. Disease-associated astrocyte epigenetic memory promotes CNS pathology. Nature 627, 865–872 (2024).

19. Park, H. et al. Single-cell RNA-sequencing identifies disease-associated oligodendrocytes in male APP NL-G-F and 5XFAD mice. Nat. Commun. 14, 802 (2023).

20. Kenigsbuch, M. et al. A shared disease-associated oligodendrocyte signature among multiple CNS pathologies. Nat. Neurosci. 25, 876–886 (2022).

21. Bohlen, C. J., Friedman, B. A., Dejanovic, B. & Sheng, M. Microglia in Brain Development, Homeostasis, and Neurodegeneration. Annu. Rev. Genet. 53, 1–26 (2019).

22. Cheng, F., Xu, J., Pieper, A. A., Cummings, J. L. & Leverenz, J. B. Discovery of disease-associated, tau, and inflammatory microglial subpopulations and molecular drivers in Alzheimer’s disease using network-based deep learning integration of human brain single-cell RNA-sequencing data. Alzheimer’s Dement. 19, (2023).

23. Takahashi, K. Microglial heterogeneity in amyotrophic lateral sclerosis. J. Neuropathol. Exp. Neurol. 82, 140–149 (2022).

24. Gao, T. et al. Transcriptional regulation of homeostatic and disease-associated-microglial genes by IRF1, LXRβ, and CEBPα. Glia 67, 1958–1975 (2019).

25. Pandey, S. et al. Disease-associated oligodendrocyte responses across neurodegenerative diseases. Cell Rep. 40, **11**1189 (2022).

26. Caglayan, E., Liu, Y. & Konopka, G. *Neuron*al ambient RNA contamination causes misinterpreted and masked cell types in brain single-nuclei datasets. Neuron 110, 4043–4056.e5 (2022).

27. Zhang, Y. et al. Ambient RNAs removal of cortex-specific snRNA-seq reveals Apoe+ microglia/macrophage after deeper cerebral hypoperfusion in mice. J. Neuroinflammation 20, **15**2 (2023).

28. Fleming, S. J. et al. Unsupervised removal of systematic background noise from droplet-based single-cell experiments using CellBender. Nat. Methods 20, 1323–1335 (2023).

29. Germain, P.-L., Lun, A., Meixide, C. G., Macnair, W. & Robinson, M. D. Doublet identification in single-cell sequencing data using scDblFinder. F1000Research 10, 979 (2021).

30. Lee, S.-H. et al. TREM2-independent oligodendrocyte, astrocyte, and T cell responses to tau and amyloid pathology in mouse models of Alzheimer disease. Cell Rep. 37, 110158 (2021).

31. Ozmen, L., Albientz, A., Czech, C. & Jacobsen, H. Expression of Transgenic APP mRNA Is the Key Determinant for Beta-Amyloid Deposition in PS2APP Transgenic Mice. Neurodegener. Dis. 6, 29–36 (2008).

32. Richards, J. G. et al. PS2APP Transgenic Mice, Coexpressing hPS2mut and hAPPswe, Show Age-Related Cognitive Deficits Associated with Discrete Brain Amyloid Deposition and Inflammation. J. Neurosci. 23, 8989–9003 (2003).

33. Jawhar, S., Trawicka, A., Jenneckens, C., Bayer, T. A. & Wirths, O. Motor deficits, neuron loss, and reduced anxiety coinciding with axonal degeneration and intraneuronal Aβ aggregation in the 5XFAD mouse model of Alzheimer’s disease. Neurobiol. Aging 33, 196.e29–196.e40 (2012).

34. Richard, B. C. et al. Gene Dosage Dependent Aggravation of the Neurological Phenotype in the 5XFAD Mouse Model of Alzheimer’s Disease. J. Alzheimer’s Dis. 45, 1223–1236 (2015).

35. Yoshiyama, Y. et al. Synapse Loss and Microglial Activation Precede Tangles in a P301S Tauopathy Mouse Model. Neuron 54, 343–344 (2007).

36. Götz, J., Chen, F., van Dorpe, J. & Nitsch, R. M. Formation of neurofibrillary tangles in P301l tau transgenic mice induced by Abeta **42** fibrils. Sci. (N. York, NY) 293, 1491–5 (2001).

37. Lee, S.-H. et al. Trem2 restrains the enhancement of tau accumulation and neurodegeneration by β-amyloid pathology. Neuron 109, 1283–1301.e6 (2021).

38. Grueninger, F. et al. Phosphorylation of Tau at S422 is enhanced by Abeta in TauPS2APP triple transgenic mice. Neurobiol. Dis. 37, 294–306 (2009).

39. Miyamura, S., Matsuo, N., Nagayasu, K., Shirakawa, H. & Kaneko, S. Myelin Oligodendrocyte Glycoprotein 35-55 (MOG 35-55)-induced Experimental Autoimmune Encephalomyelitis: A Model of Chronic Multiple Sclerosis. BIO-Protoc. 9, e3453 (2019).

40. Wheeler, M. A. et al. MAFG-driven astrocytes promote CNS inflammation. Nature 578, 593–599 (2020).

41. DerSimonian, R. & Laird, N. Meta-analysis in clinical trials. Control. Clin. Trials 7, 177–188 (1986).

42. Wang, L. et al. Primary cilia signaling in astrocytes mediates development and regional-specific functional specification. Nat. Neurosci. 27, 1708–1720 (2024).

43. Horan, K. & Williams, A. C. Mapping out multiple sclerosis with spatial transcriptomics. Nat. Neurosci. 27, 2270–2272 (2024).

44. Hao, Y. et al. Integrated analysis of multimodal single-cell data. Cell 184, 3573–3587.e29 (2021).

45. Farhy-Tselnicker, I. et al. Activity-dependent modulation of synapse-regulating genes in astrocytes. eLife 10, e70514 (2021).

46. Tewari, B. P. et al. Astrocytes require perineuronal nets to maintain synaptic homeostasis in mice. Nat. Neurosci. 27, 1475–1488 (2024).

47. Wilkins, H. M. & Swerdlow, R. H. Mitochondrial links between brain aging and Alzheimer’s disease. Transl. Neurodegener. 10, 33 (2021).

48. Chandler, H. L. et al. Reduced brain oxygen metabolism in patients with multiple sclerosis: Evidence from dual-calibrated functional MRI. J. Cereb. Blood Flow Metab. 43, 115–128 (2022).

49. Hasel, P., Rose, I. V. L., Sadick, J. S., Kim, R. D. & Liddelow, S. A. Neuroinflammatory astrocyte subtypes in the mouse brain. Nat. Neurosci. 24, 1475–1487 (2021).

50. Rao, S., Yung, J., Edick, M. G. & Hanson, J. E. Protocol for detection of glial complement expression in relation to amyloid plaques in mouse brain with combined FISH and IHC. STAR Protoc. 5, 103388 (2024).

51. Batiuk, M. Y. et al. Identification of region-specific astrocyte subtypes at single cell resolution. Nat. Commun. 11, 1220 (2020).

52. Fleming, S. J. et al. Unsupervised removal of systematic background noise from droplet-based single-cell experiments using CellBender. Nat. Methods 20, 1323–1335 (2023).

53. Smajić, S. et al. Single-cell sequencing of human midbrain reveals glial activation and a Parkinson-specific neuronal state. Brain awab446- (2021) doi:10.1093/brain/awab446.

54. Jäkel, S. et al. Altered human oligodendrocyte heterogeneity in multiple sclerosis. Nature 566, 543–547 (2019).

55. Schirmer, L. et al. Neuronal vulnerability and multilineage diversity in multiple sclerosis. Nature 573, 75–82 (2019).

56. Morabito, S. et al. Single-nucleus chromatin accessibility and transcriptomic characterization of Alzheimer’s disease. Nat Genet 53, 1143–1155 (2021).

57. Sadick, J. S. et al. Astrocytes and oligodendrocytes undergo subtype-specific transcriptional changes in Alzheimer’s disease. Neuron 110, 1788–1805.e10 (2022).

58. Macnair, W., et al. Single nuclei RNAseq stratifies multiple sclerosis patients into distinct white matter glial responses. bioRxiv 2022.04.06.487263 (2023) doi:10.1101/2022.04.06.487263.

59. Wang, Q. et al. Single-cell transcriptomic atlas of the human substantia nigra in Parkinson’s disease. bioRxiv 2022.03.25.485846 (2022) doi:10.1101/2022.03.25.485846.

60. Gabitto, M. I., et al. Integrated multimodal cell atlas of Alzheimer’s disease. bioRxiv 2023.05.08.539485 (2024) doi:10.1101/2023.05.08.539485.

61. Cain, A. et al. Multicellular communities are perturbed in the aging human brain and Alzheimer’s disease. Nat. Neurosci. 26, 1267–1280 (2023).

62. Absinta, M. et al. A lymphocyte–microglia–astrocyte axis in chronic active multiple sclerosis. Nature 597, 709–714 (2021).

63. Kamath, T. et al. Single-cell genomic profiling of human dopamine neurons identifies a population that selectively degenerates in Parkinson’s disease. Nat. Neurosci. 25, 588–595 (2022).

64. Gerrits, E. et al. Distinct amyloid-β and tau-associated microglia profiles in Alzheimer’s disease. Acta Neuropathol. 141, 681–696 (2021).

65. Smith, A. M. et al. Diverse human astrocyte and microglial transcriptional responses to Alzheimer’s pathology. Acta Neuropathol. 143, 75–91 (2022).

66. Korsunsky, I. et al. Fast, sensitive, and accurate integration of single cell data with Harmony. Nat. methods 16, 1289–1296 (2019).

67. Hodge, R. D. et al. Conserved cell types with divergent features in human versus mouse cortex. Nature 573, 61–68 (2019).

68. Irwin, A. B. et al. The lncRNA Neat1 is associated with astrocyte reactivity and memory deficits in a mouse model of Alzheimer’s disease. bioRxiv 2023.05.03.539260 (2023) doi:10.1101/2023.05.03.539260.

69. Journiac, N. et al. The nuclear receptor RORα exerts a bi-directional regulation of IL-6 in resting and reactive astrocytes. Proc. Natl. Acad. Sci. 106, 21365–21370 (2009).

70. Liu, Z. et al. Astrocytic response mediated by the CLU risk allele inhibits OPC proliferation and myelination in a human iPSC model. Cell Rep. 42, 112841 (2023).

71. Yu, W. et al. Disease-Associated Neurotoxic Astrocyte Markers in Alzheimer Disease Based on Integrative Single-Nucleus RNA Sequencing. Cell. Mol. Neurobiol. 44, 20 (2024).

72. Wang, W., Lv, R., Zhang, J. & Liu, Y. circSAMD4A participates in the apoptosis and autophagy of dopaminergic neurons via the miR-29c-3p-mediated AMPK/mTOR pathway in Parkinson’s disease. Mol. Med. Rep. 24, 540 (2021).

73. Anderson, A. G. et al. Single nucleus multiomics identifies ZEB1 and MAFB as candidate regulators of Alzheimer’s disease-specific cis-regulatory elements. Cell Genom. 3, 100263 (2023).

74. Li, Z.-D. et al. The divergent effects of astrocyte ceruloplasmin on learning and memory function in young and old mice. Cell Death Dis. 13, 1006 (2022).

75. Kim, R. D. et al. Temporal and spatial analysis of astrocytes following stroke identifies novel drivers of reactivity. bioRxiv 2023.11.12.566710 (2023) doi:10.1101/2023.11.12.566710.

76. Liddelow, S. A. et al. Neurotoxic reactive astrocytes are induced by activated microglia. Nature 541, 481–487 (2017).

77. Shi, Q. et al. Complement C3 deficiency protects against neurodegeneration in aged plaque-rich APP/PS1 mice. Sci. Transl. Med. 9, (2017).

78. Wu, T. et al. Complement C3 Is Activated in Human AD Brain and Is Required for Neurodegeneration in Mouse Models of Amyloidosis and Tauopathy. Cell Rep. 28, 2111–2123.e6 (2019).

79. Lindia, J. A., McGowan, E., Jochnowitz, N. & Abbadie, C. Induction of CX3CL1 Expression in Astrocytes and CX3CR1 in Microglia in the Spinal Cord of a Rat Model of Neuropathic Pain. J. Pain 6, 434–438 (2005).

80. Asano, S., et al. Microglia–Astrocyte Communication via C1q Contributes to Orofacial Neuropathic Pain Associated with Infraorbital Nerve Injury. Int. J. Mol. Sci. 21, 6834 (2020).

81. Catalano, M. et al. CX3CL1 protects neurons against excitotoxicity enhancing GLT-1 activity on astrocytes. J. Neuroimmunol. 263, 75–82 (2013).

82. Cheli, V. T. et al. The expression of ceruloplasmin in astrocytes is essential for postnatal myelination and myelin maintenance in the adult brain. Glia 71, 2323–2342 (2023).

83. Rothhammer, V. et al. Sphingosine 1-phosphate receptor modulation suppresses pathogenic astrocyte activation and chronic progressive CNS inflammation. Proc. Natl. Acad. Sci. 114, 2012–2017 (2017).

84. Dozio, V. & Sanchez, J.-C. Profiling the proteomic inflammatory state of human astrocytes using DIA mass spectrometry. J. Neuroinflammation 15, 331 (2018).

85. Simon, C. M. et al. Dysregulated IGFBP5 expression causes axon degeneration and motoneuron loss in diabetic neuropathy. Acta Neuropathol. 130, 373–387 (2015).

86. Rauskolb, S. et al. Insulin-like growth factor 5 associates with human Aß plaques and promotes cognitive impairment. Acta Neuropathol. Commun. 10, 68 (2022).

87. Crow, M., Paul, A., Ballouz, S., Huang, Z. J. & Gillis, J. Characterizing the replicability of cell types defined by single cell RNA-sequencing data using MetaNeighbor. Nat. Commun. 9, 884 (2018).

88. Jung, M. et al. Cross-species transcriptomic atlas of dorsal root ganglia reveals species-specific programs for sensory function. Nat. Commun. 14, 366 (2023).

89. Ling, E. et al. A concerted neuron–astrocyte program declines in ageing and schizophrenia. Nature 627, 604–611 (2024).

90. Gabitto, M. I. et al. Integrated multimodal cell atlas of Alzheimer’s disease. Nat. Neurosci. 27, 2366–2383 (2024).

91. Green, G. S. et al. Cellular communities reveal trajectories of brain ageing and Alzheimer’s disease. Nature 633, 634–645 (2024).

92. Serrano-Pozo, A. et al. Astrocyte transcriptomic changes along the spatiotemporal progression of Alzheimer’s disease. Nat. Neurosci. 27, 2384–2400 (2024).

93. Wu, T. et al. clusterProfiler 4.0: A universal enrichment tool for interpreting omics data. Innov. 2, 100141 (2021).

94. Subramanian, A. et al. Gene set enrichment analysis: A knowledge-based approach for interpreting genome-wide expression profiles. Proc. Natl. Acad. Sci. 102, 15545–15550 (2005).

