## Supplementary material for "Integrated Cross-Disease Atlas of Human And Mouse Astrocytes Reveals Heterogeneity and Conservation of Astrocyte Subtypes in Neurodegeneration": All Supplemental Figures

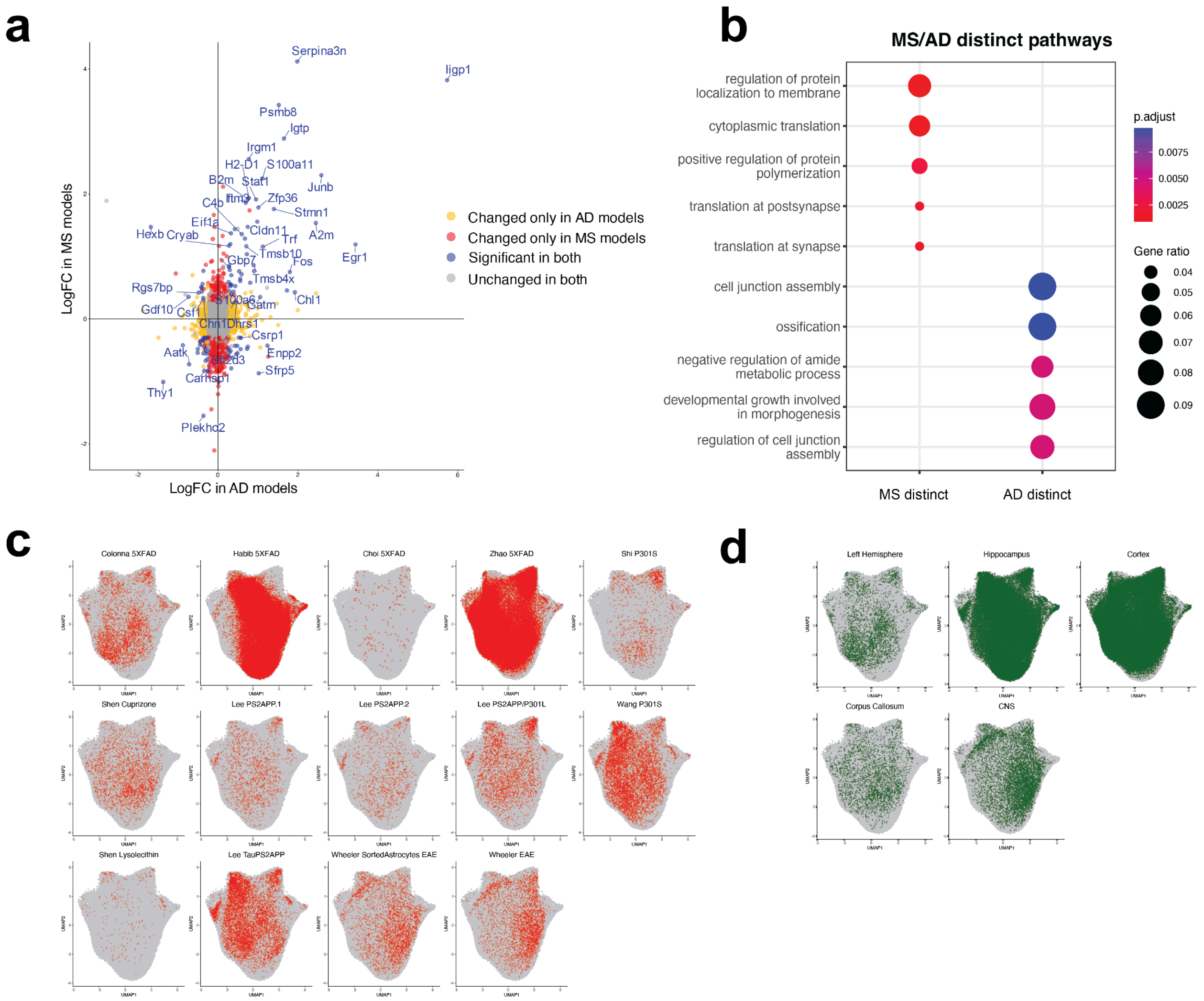


***Supplemental Figure 1.***

*(A) Comparison of differentially expressed genes between mouse astrocytes AD vs. control and MS vs. control. The y-axis represents log-fold changes in mouse AD models vs. controls, and the x-axis represents log-fold changes in mouse MS models vs controls. Color represents significantly changed genes in both diseases (blue) only in MS (red) only in AD (yellow) or not changed in either (gray). (B) Dotplot of Gene Ontology overrepresentation analysis of shared and distinct differentially expressed genes in AD and MS. The color of the circles indicates the adjusted p-value, and the size indicates the ratio of genes in that pathway. Uniform Manifold Approximation and Projection (UMAP) of mouse neurodegeneration and control astrocytes, faceted by study(C) and by brain region (D).*

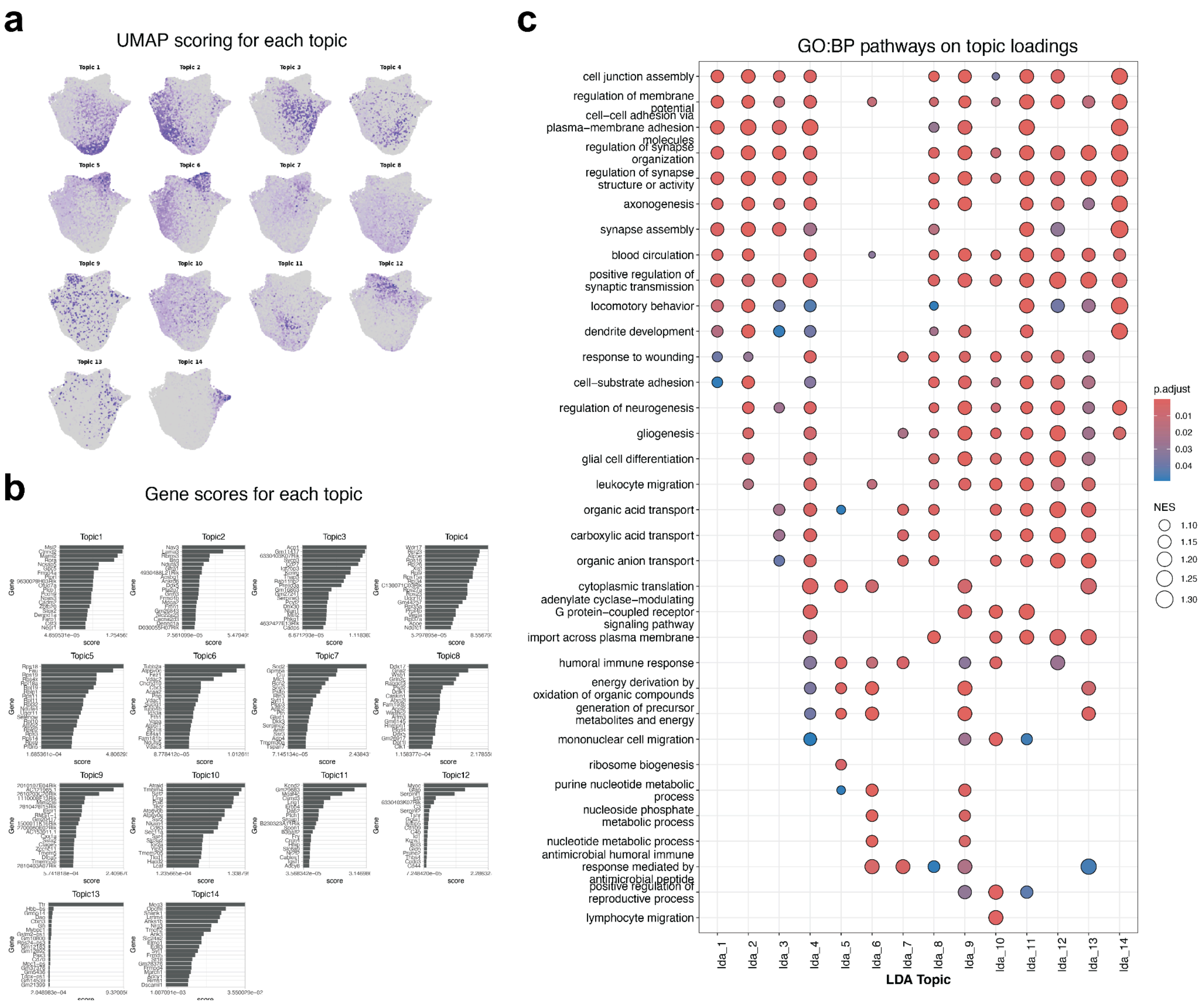


***Supplemental Figure 2.***

(A) *Uniform Manifold Approximation and Projection (UMAP) of mouse neurodegeneration and control astrocytes scoring for topic loadings. (B) Barplots depicting the top 10 genes defining the topic. (C) Dotplot of Gene Ontology overrepresentation analysis of genes defining topics. The color of the circles indicates the adjusted p-value, and the size indicates the ratio of genes in that pathway.*

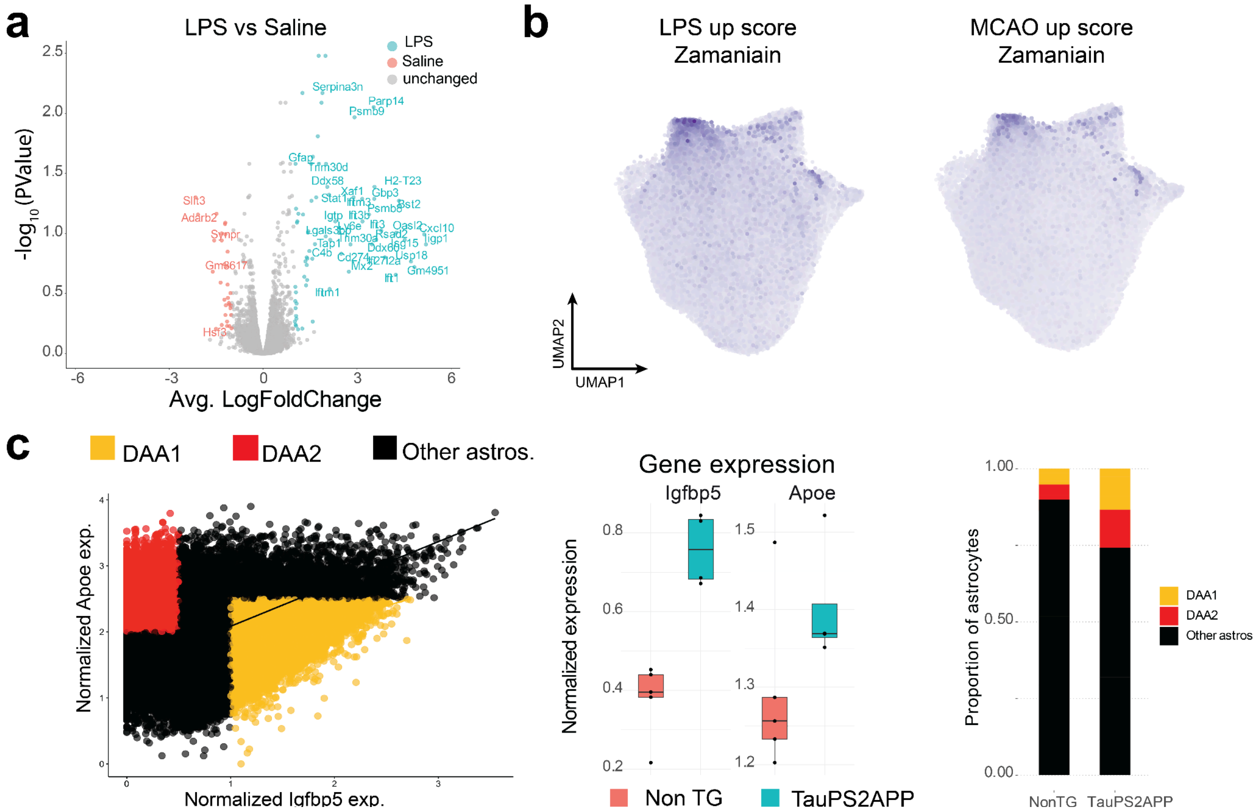


***Supplemental Figure 3.***

(A)*Volcano Plot indicating differential expression between LPS and Saline astrocytes. Orange and blue colors indicate significantly differentially expressed genes (meta-logfoldChange > 0.5, metaFDR < 0.05), and gray indicates no significant change (See Methods). (B) Scoring mouse-integrated neurodegeneration astrocytes with differentially expressed genes from LPS and MCAO astrocytes. UMAP shows the distribution of expression scores and enrichment in the DAA1 population. (C)* In-situ RNA analysis of astrocyte subtypes depicted in Figure 4. Plot shows normalized Apoe expression vs. normalized Igfb5 expression among all cells with high Slc1a3 expression. Expression was normalized to cell size. Normalized expression cutoffs for DAA1 (yellow; moderate Apoe and high Igfbp5) and DAA2 (red; high ApoE and low Igfbp5) are shown, with all other Slc1a3 high cells categorized as Other Astrocytes (black) (n = 55,492 astrocytes from 2 Non-TG and 4 TauPS2APP mice) (D) Compared to non-transgenic control mice (Non-TG), TauPS2APP mice had higher levels of expression of Igfbp5 and Apoe (left) and correspondingly exhibited a higher average proportion of DAA1 and DAA2 cells (n = 2 Non TG and 4 TAuPS2APP) *(right).*

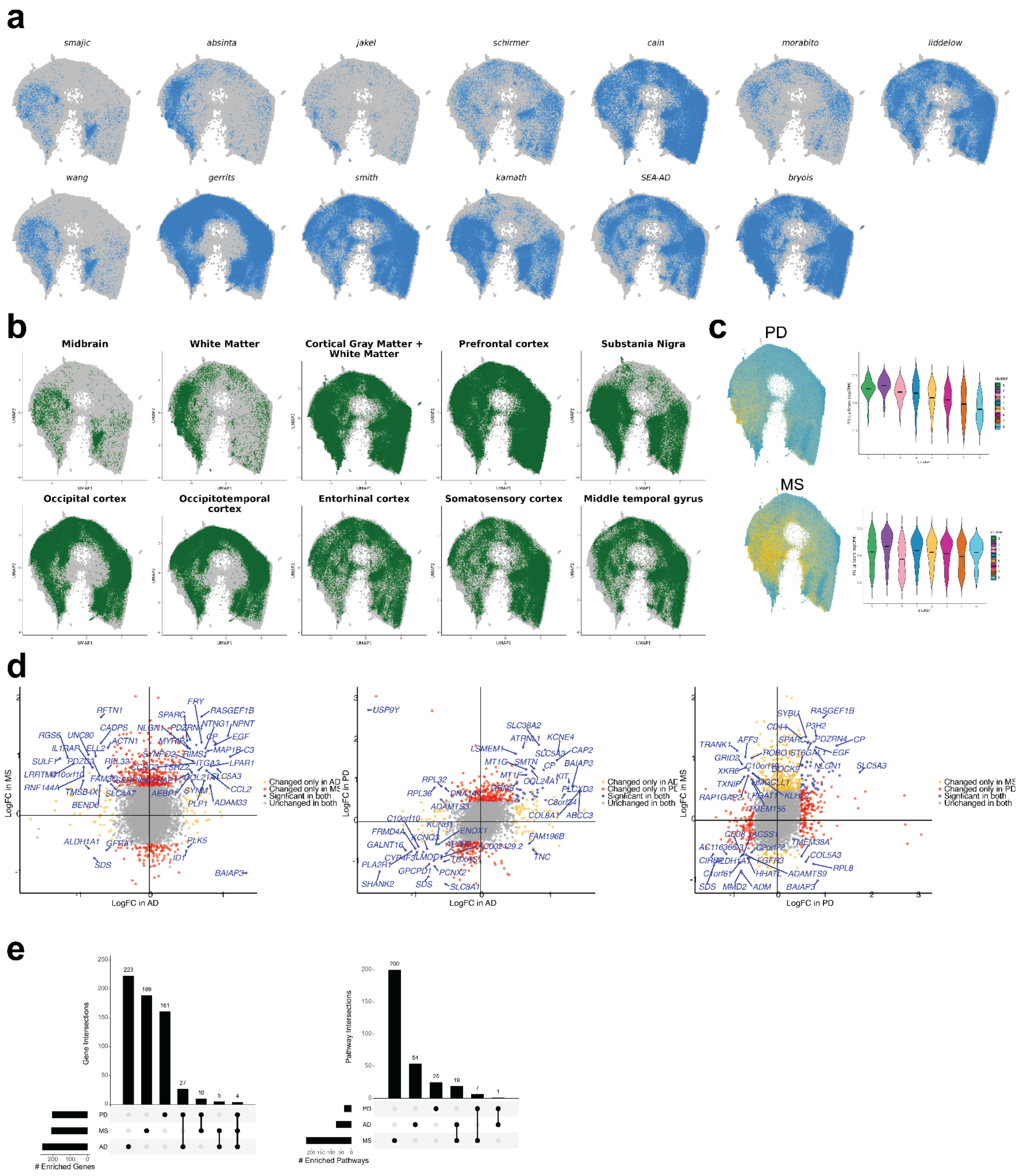


### *Supplemental Figure 4*

*Uniform Manifold Approximation and Projection (UMAP) of human neurodegeneration and control astrocytes, faceted by study(A) and by brain region (B). (C) Scoring human integrated neurodegeneration astrocytes with differentially expressed genes from MS and PD astrocytes. UMAP shows the distribution of expression scores and enrichment in the GFAP-hi population. (D) Comparison of differentially expressed genes between human astrocytes AD vs. control and MS vs. control (left), AD vs. control and PD vs. control (middle), and PD vs. control and MS vs. control (right). Color represents significantly changed genes in both species (blue) only in disease 1 (red) and disease 2 (yellow), or they are not changed in either disease (gray). (E) Upset Plot depicting shared and distinct gene expression changes across all three diseases.*

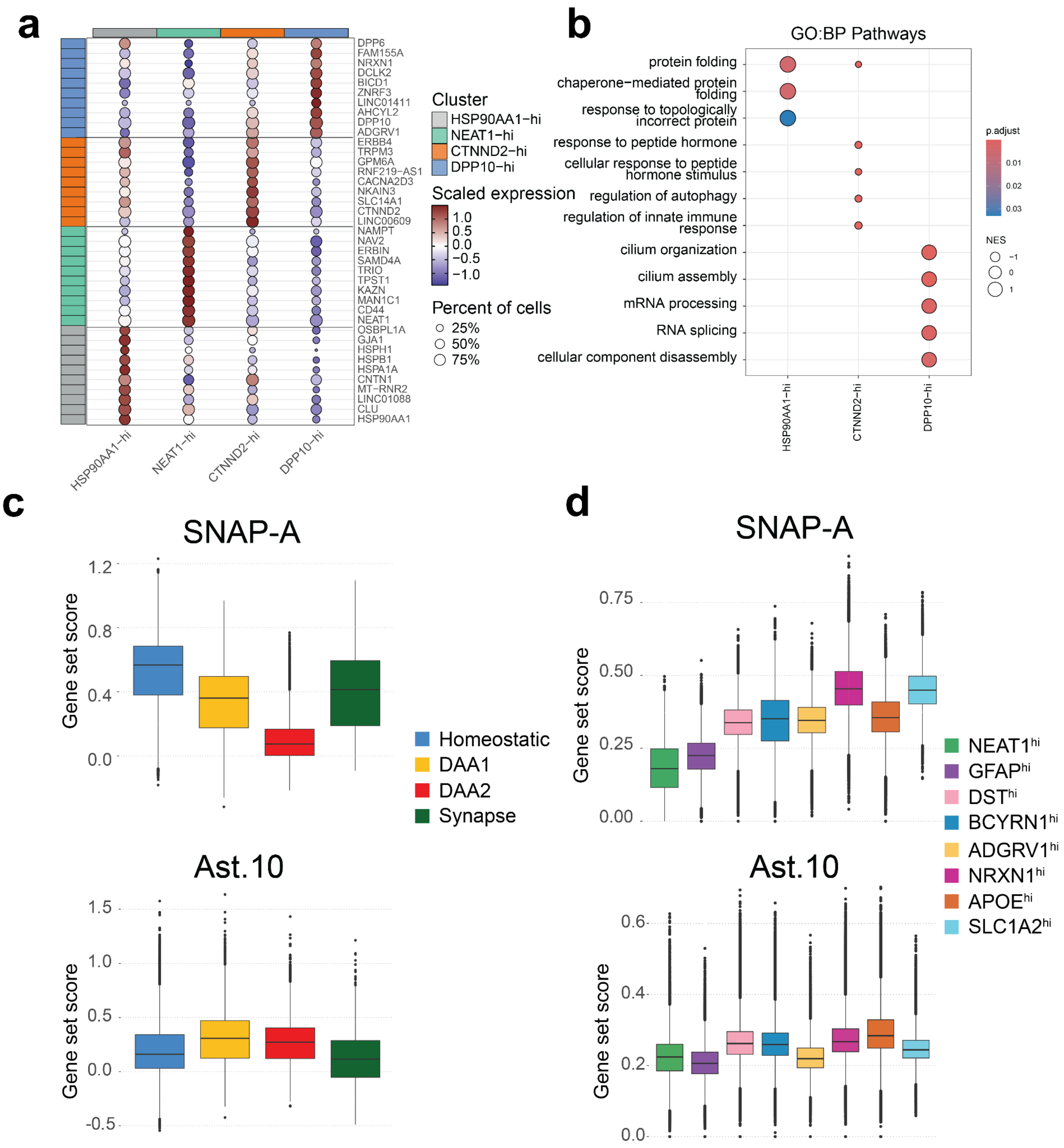


***Supplemental Figure 5.***

*(A) Dotplot of the top 5 cluster markers of each GFAP-hi human subcluster. Color represents the scaled expression of each cell in that cluster. Dot size represents the percentage of cells that express that specific marker. (B) Dotplot of top differential*

*pathways. The color of the circles indicates the adjusted p-value, and the size indicates the Normalized Enrichment Score (NES) for the specific pathway. Row names include the gene ontology categories. Pathways were identified using Gene Set Enrichment analysis on Gene Ontology biological pathways. (C) Scoring integrated neurodegeneration astrocytes shown in Figure 2A with SNAP-A gene program (top). Screenshot for our searchable website of top gene in SNAP-A program Gpc5, shown in both mouse (middle) and human (bottom). (D) Screenshot of cluster expression from our searchable website for gene of interest MT2 in both human and mouse.*
